# Step-wise activation of a Family C GPCR

**DOI:** 10.1101/2023.08.29.555158

**Authors:** Kaavya Krishna Kumar, Haoqing Wang, Chris Habrian, Naomi R. Latorraca, Jun Xu, Evan S. O’Brien, Chensong Zhang, Elizabeth Montabana, Antoine Koehl, Susan Marqusee, Ehud Y. Isacoff, Brian K. Kobilka

## Abstract

Metabotropic glutamate receptors belong to a family of G protein-coupled receptors that are obligate dimers and possess a large extracellular ligand-binding domain (ECD) that is linked via a cysteine-rich domain (CRDs) to their 7-transmembrane (TM) domain. Upon activation, these receptors undergo a large conformational change to transmit the ligand binding signal from the ECD to the G protein-coupling TM. In this manuscript, we propose a model for a sequential, multistep activation mechanism of metabotropic glutamate receptor subtype 5. We present a series of structures in lipid nanodiscs, from inactive to fully active, including agonist-bound intermediate states. Further, using bulk and single-molecule fluorescence imaging we reveal distinct receptor conformations upon allosteric modulator and G protein binding.

## Introduction

Metabotropic glutamate receptors (mGlus) belong to a family of obligate dimeric G protein-coupled receptors (GPCRs), that are activated by the excitatory neurotransmitter, *L*-glutamate^1^. Each protomer contains a large ECD that is made up of a Venus fly trap (VFT) domain, that contains the orthosteric ligand binding site, and a cysteine-rich domain (CRD). The CRD connects the VFT to the family-defining 7-transmembrane (TM) domain (Extended Data Figure 1a)^1^. The binding of glutamate causes the closure of the VFTs and a protomer rearrangement that brings the CRDs and TM domains in close proximity^2^. Dimerization of the mGlus is mandatory for their function, and the rearrangement upon activation of the receptors is a complex allosteric process with the two protomers influencing each other^3^. Allosteric modulators of mGlus bind the TM domain^4^ to regulate signaling by themselves or in conjunction with orthosteric ligands. Ultimately, agonists and positive allosteric modulators (PAMs) activate mGlus by stabilizing intermolecular interactions between the protomers which enable G protein coupling to the TM domain.

To better understand the activation mechanism of mGlus, and to delineate the allosteric link between ligands and G proteins, we took mGlu5 as a representative of the mGlu family and employed a combination of structural and biophysical techniques to characterize the conformational landscape of receptor activation. We first show the importance of a lipid bilayer for mGlu5 activation. To comprehensively understand the receptor activation pathway, we determine structures of mGlu5 in the presence of a small molecule orthosteric agonist and an allosteric modulator, as well as the allosteric nanobody, Nb43^2^. As a pure PAM, Nb43 potentiates the activity of orthosteric agonists but lacks intrinsic activity of its own. In the presence of the orthosteric agonist L-quisqualic acid (Quis) and Nb43, we resolved two receptor conformations; (i) an intermediate (Intermediate 1a, Figure 1a) with the VFT upper lobes closed but a large interprotomer distance reflecting an inactive conformation, and (ii) an “active-like” structure (Intermediate 2a, Figure 1a) where the VFT lower lobes, CRDs and the TMs are in close proximity. It should be noted that for the purpose of discussion, all conformations between Apo and G protein-bound “Fully Active”, are named “Intermediates”. Addition of agonist-PAM 3-cyano-*N*-1,3-diphenyl-1*H*-pyrazol-5-yl)benzamide (CDPPB) to the Quis- and Nb43-bound mGlu5 fully stabilized a single active conformation (Intermediate 3a, Figure 1a) that resembles the Quis and Nb43-bound Intermediate 2a structure. Interestingly, in this complex, we observe an asymmetric action of CDPPB, with density supporting its binding to only one of the two TMs, as opposed to a structure of mGlu5 with CDPPB alone, but *without* Quis or Nb43, where we observe CDPPB in both TMs (Intermediate 1b, Figure 1a). We further investigate the intersubunit conformational changes upon ligand and G protein binding using bulk fluorescence spectroscopy and single-molecule Förster resonance energy transfer (smFRET). We observe receptor conformations that correlate well with the determined structures as well as several receptor conformations that are uniquely observed in biophysical studies, including states stabilized by CDPPB (Intermediates 2b and 3b, Figure 1a) and one stabilized by G protein (fully active). Finally, combining all the data we propose a model for the step-wise activation of mGlu5. Figure 1a gives an overview of the model developed from the results presented in this manuscript.

**Fig 1:**
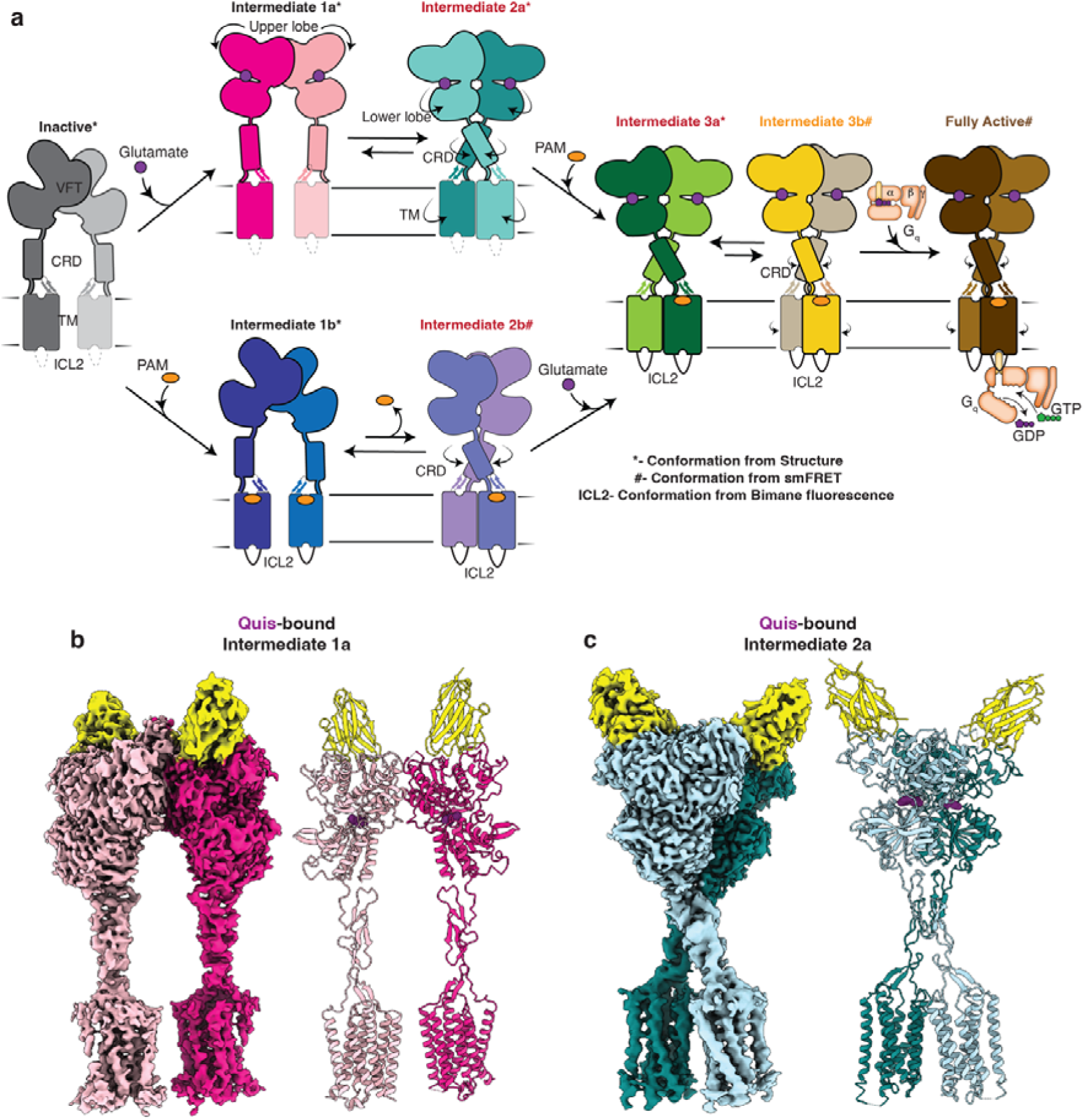
Sequential activation of mGlu5 in lipid environment. a) Using the data from this study we propose a model for mGlu5 activation. The addition of an orthosteric agonist (e.g. glutamate) results in the closing of the upper lobe (Intermediate 1a). This conformation is in equilibrium with a conformation in which the twisting of the lower lobe brings the CRDs and TMs in close proximity (Intermediate 2a). The addition of a PAM stabilizes the CRDs and TMs, including ICL2 in an active conformation (Intermediate 3a). Intermediate 3a is in equilibrium with Intermediate 3b, which is characterized by a smaller intersubunit distance. In the presence of an orthosteric agonist, the PAM binds to one protomer (Intermediate 3a and 3b), whereas in its absence the PAM binds to both the protomers symmetrically (Intermediate 1b). Further addition of G protein to the agonist and PAM-bound mGlu5 results in the stabilization of a unique fully active conformation of the receptor (Fully active). b) Cryo-EM density and model of Quis-bound mGlu5 in nanodisc, representing an Intermediate 1a state, where Quis is bound to the VFTs, however, the CRDs and TMs are far apart mimicking the inactive state. VFT binding Nb43 is shown in yellow. c) Cryo-EM density and model of nanodisc-incorporated Quis-bound mGlu5, Intermediate 2a state. The CRDs and TMs are in an active conformation (close together).

## Results

### A lipid environment is critical for mGlu5 activation

To identify the best conditions for studying mGlu5 activation, we monitored the ability of mGlu5 to drive GTP turnover via the heterotrimeric G protein G_q_ in detergent micelles and reconstituted in lipid nanodiscs using the belt protein MSP2N2 (~ 16 nm diameter)^5^ and a POPC/POPG lipid mixture (Extended Data Figure 1b). In detergent micelles, we observed no activation of G_q_ by mGlu5 bound to Quis, and only minimal activation (~ 20 % above G_q_ alone) by mGlu5 bound to both Quis and CDPPB (Extended Data Figure 1c), while the M1 muscarinic receptor in detergent micelles was able to efficiently activate G_q_ (Extended Data Figure 1d). In contrast, mGlu5 in lipid nanodiscs was able to robustly activate G_q_ by Quis alone or Quis and CDPPB (Extended Data Figure 1c). To assess whether this dependence on lipid environment for mGlu5 activation is a consequence of alteration in receptor conformation, rather than a direct effect of lipid on either G_q_ activation or mGlu5 affinity for ligand, we performed hydrogen-deuterium exchange monitored by mass spectrometry (HDX-MS). Upon agonist binding to mGlu5 in detergent, we observe changes in the VFT, consistent with ligand binding^2^ (Extended Data Figure 1e). These VFT changes in detergent are nearly identical to HDX-MS curves of agonist-bound mGlu5 in nanodiscs (Extended Data Figure 1e-f). Thus, the effects of agonist binding on the VFT do not appear to depend on the receptor TM environment. However, there are notable differences in the TM region of agonist-bound mGlu5 between detergent and nanodiscs: specifically, peptides in the intracellular region of TM3 exhibit reduced deuterium uptake in mGlu5 in nanodiscs compared to mGlu5 in detergents (Extended Data Figure 1e, g). Other peptides in the TM region, including in TM5, do not exhibit differences in deuterium uptake, suggesting that the detergent environment does not globally destabilize the receptor (Extended Data Figure 1g). In other mGlus, e.g. in mGlu2, this region of TM3 undergoes conformational changes upon activation and interacts with G protein^6^ (Extended Data Figure 1h). Taken together, these data are consistent with a model where lipids modulate the ability of mGlu5 to adopt an active state capable of GTP turnover.

### Quis-bound mGlu5 exists in two conformation

To better understand the activation mechanism of mGlu5, we determined the structure of Quis and Nb43 bound mGlu5 reconstituted into lipid nanodisc using single particle cryo-electron microscopy (cryo-EM). Data processing revealed two distinct conformations of the receptor (Extended Data Figure 2). Though both structures have Quis bound (to both protomers), one shows an “intermediate” conformation (Intermediate 1a, Figure 1a) wherein the VFTs are closed, but CRDs and TMs are far apart as in the inactive mGlu5 structure (Figures 1b, Extended Figure 3a), while the other conformation is “active-like” (Intermediate 2a, Figure 1a) such that the CRDs and TMs are close together forming the TM6-TM6 interface, the hallmark of Family C activation^2^ (Figure 1c).

To investigate changes accompanying Quis binding, we compared the previously determined Apo (PDB code: 6N52^2^) structure to the current Quis-bound, Intermediate 1a structure (Extended Data Figure 3a). Upon Quis binding, there is a decrease in distance between the upper and lower lobes of the VFT (Figure 2a). This decrease is mostly due to the closure of the upper lobe (Figure 2a, Extended Data Figure 3b). The upper lobe closure is accompanied by some rearrangements in the lower lobe and the “hinge region” (the region between the lobes in the dimer interface (Extended Data Figure 1a, Extended Data Figure 3b). For example, changes to Quis interacting residues such as W100 and E279 (Figure 2b, Extended Data Figure 3a) result in conformational changes in the B and C helices at the intersubunit interface (Figure 2a-b). Hindrance to the movement of these residues appears to stabilize the receptor in an inactive conformation, as seen in the antagonist-bound structure (PDB code: 7FD9^7^) (Extended Data Figure 3c). Hence, changes in the lower lobe residues (e.g. E279) and the “hinge region” triggers rearrangement of the upper lobes (i.e. closing) upon Quis binding (Figure 2a, Extended Data Figure 3b).

**Fig 2:**
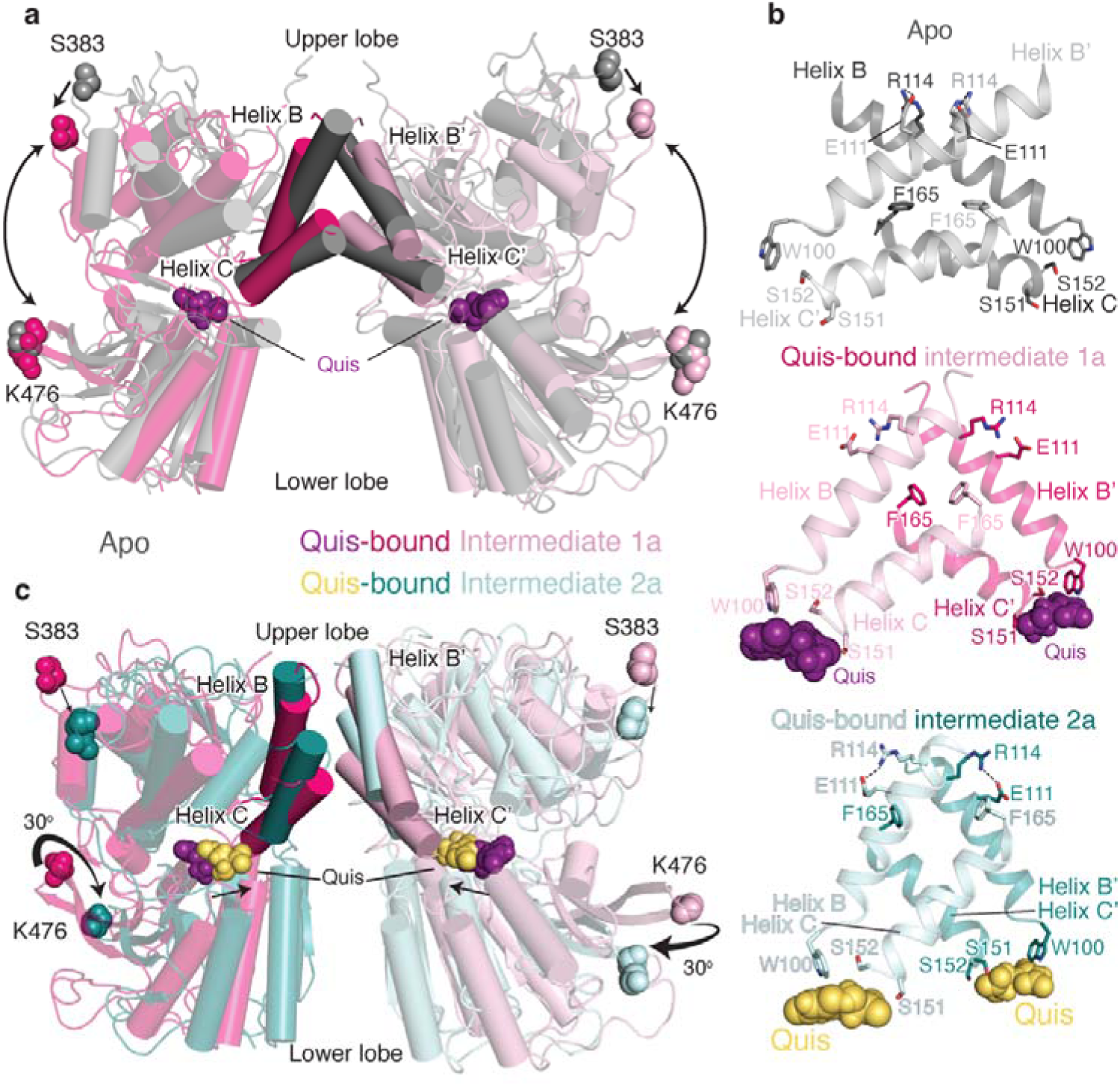
Structures of Quis-bound conformations of mGlu5 in nanodisc. a) VFTs of Apo (grey, PDB: 6N52) and Quis-bound Intermediate 1a are overlayed. Upon Quis binding the upper lobe closes, as seen by the movement of S383, whereas not much change is seen in the lower lobe (comparing K476 between the structures). Also shown is the comparison of the B and C helices at the intersubunit interface in the Apo and Quis-bound Intermediate 1a state. b) The intersubunit rearrangement upon Quis binding reorients the B and C helices leading to a reduction in the helix angle. Top: Apo, Middle: Quis-bound Intermediate 1a and Bottom: Quis-bound Intermediate 2a. Residue R114 interacts with E111 from the adjacent protomer in the Apo state and within the same protomer in the Quis-bound Intermediate 2a. The residue F165 is shown to illustrate the change in the position of the C helix. There is a downward movement of W100 towards Quis in Intermediates 1a and 2a. Due to the lower lobe rotation in Intermediate 2a, a further inward movement of Quis is seen. c) Overlay of VFTs of Quis-bound Intermediate 1a and Quis-bound Intermediate 2a showing a small change in the upper lobe (movement of S383). The lower lobes twist 30° and move closer together as seen by comparing K476 between the structures. The B and C helices at the protomer-protomer interface in the Quis-bound Intermediate 2a state show an upward shift compared to the Quis-bound Intermediate 1a. This likely is the result of the inward movement of Quis (from purple to yellow) and the rearrangement of the lower lobe.

To study the structural changes that occur subsequent to Quis binding to form the active state of the receptor, we compared the Quis-bound Intermediate 1a (Figure 1a) and the Quis-bound Intermediate 2a (Figure 1a) structures of mGlu5 (Figure 2b-c). We observe a large “twisting” of the VFT lower lobe in the Intermediate 2a structure(Figure 2c, Extended Data Figure 4a). The lower lobe of the VFT moves as a rigid body and maintains the Quis binding pocket (RMSD ~ 0.5, Extended Data Figure 4b) and initiates the rearrangement of the B and C helices in the hinge region (Figure 2b, c). These structural changes ultimately lead to the CRDs and TMs (Extended Data Figure 5) moving close to each other (hallmark of Family C GPCR activation) to activate mGlu5.

### Symmetric to asymmetric binding of mGlu5 PAM

Allosteric modulators of Family C GPCRs bind in the TM region and modulate orthosteric ligand binding and signaling. Crystal structures of single mGlu TM domain bound to negative allosteric modulators (NAMs) have been previously determined^8–10^. To gain structural insights into the activity of allosteric modulators in the context of the full-length receptor, we determined the structures of nanodisc incorporated CDPPB-bound mGlu5 (and Nb43) in the absence and presence of Quis (Figure 3a, Extended Data Figure 6–7). In the absence of Quis, CDPPB-bound mGlu5 (Intermediate 1b, Figure 1a) adopts a conformation in which the protomers are separated and we observe density for CDPPB in both TM protomers indicating *symmetric* binding (similar to previously seen for mGlu bound to NAM ^11,12^) (Extended Data Figure 8). However, in the presence of Quis, CDPPB binds to the TM of only one protomer (i.e. is bound *asymmetrically*) (Figure 3a). Moreover, unlike Quis alone, in Quis and CDPPB we obtained only one conformation with the protomers together (Intermediate 3a, Figure 1a, Extended Data Figure 6), consistent with stabilization of the active conformation (Extended Data Figure 9a-b).

**Fig 3:**
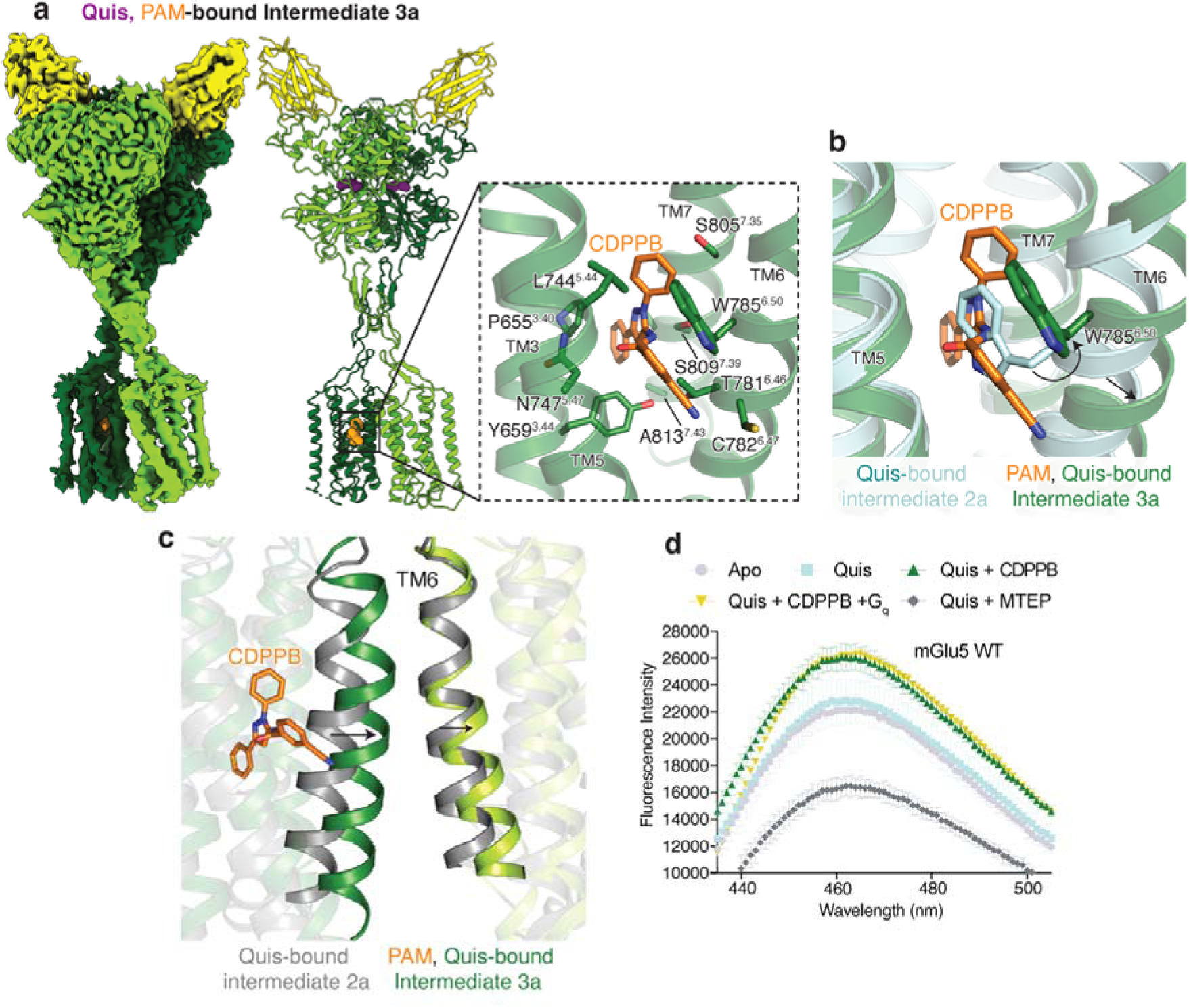
Structural changes of upon PAM binding to mGlu5. a) Cryo-EM density and model of CDPPB (orange) and Quis-bound mGlu5 in a nanodisc. The structure represents the Intermediate 3a state with the CRDs and TMs in close proximity. Nb43 is shown in yellow. Insert: Binding pocket of CDPPB in the TM region showing residues within 4Å as sticks. b) CDPPB binding to the TM causes the rearrangement of W785^6.50^ to accommodate the ligand. c) Quis-bound Intermediate 2a and CDPPB, Quis-bound Intermediate 3a structures show differences in the conformation of TM6 at the protomer interface. d) Bimane spectra of mGlu5 in nanodiscs labeled at positions C691^4.30^ (end of TM4) and C681^ICL2^. Adding Quis (cyan) results in no change in the spectra compared to Apo (grey). However, Quis and CDPPB increase the fluorescence (dark green), indicating a change in the ICL2 environment. Further addition of G_q_ does not result in a change in the bimane spectrum (yellow). The addition of Quis and MTEP causes a decrease in fluorescence. Data represented as mean ± SD, n = 3 individual.

Asymmetric allosteric modulator binding to only one protomer TM is seen in other Family C GPCRs^6^. CDPPB interacts similarly to the mGlu5 NAM, MPEP (PDB code: 6FFI^10^); with the exception of a large outward motion of W785^6.50^ (Figure 3b, Extended Data Figure 9c) (superscript indicating GPRDB canonical numbering scheme^13^). In the CDPPB-bound protomer, the entire TM6 is moved outward (C785^6.64^ Cα-Cα ~ 3.5 Å), with Y779^6.44^ pointing towards the protomer that does not contain CDBBP (Figure 3c, Extended Data Figure 9d-e). This appears to result in the inward movement of TM6 in the CDBBP-less protomer, with W785^6.50^ occluding the PAM binding site (Extended Data Figure 9f). This asymmetric PAM binding allows the protomer TMs to form a tight interface.

To tease out the differences between the Quis-bound Intermediate 2a and CDPPB, Quis-bound Intermediate 3a states, we carried out three-dimensional variability analysis (3DVA)^14^. In both cases, we observed similar modes of variability (SI Video 1), suggesting these are of biological relevance, rather than being artifacts resulting from overfitted noise in cryo-EM data. Notably, in the Quis-bound dataset, we detect a more pronounced “stretch” motion between VFT and TM domains (SI Video 1). Given that CDPPB binds the TM domain, we hypothesize that CDPPB can “lock” the TM-TM interface of mGlu5 homodimer in an active conformation, consistent with the role of CDPPB as a PAM. In addition, 3D flexible refinement (3Dflex)^15^ revealed an asymmetric stretching pattern (SI Video 2), wherein one protomer exhibited greater translational movement than the other. This indicates some asymmetry in the activation of the protomers in the presence of CDPPB, perhaps consistent with the observed asymmetric PAM binding (Figure 3a).

### Effect of ligand binding on ICL2 conformation

Previous structural studies of mGlu receptors have shown that ICL2 is stabilized in the presence of G protein^6,16^. To investigate conformational changes in ICL2 following ligand binding, we used the environmentally sensitive fluorophore, monobromobimane (bimane) as a conformational reporter (Extended Data Figure 10a-c). We performed bimane spectroscopy on two versions of nanodisc-incorporated mGlu5, the WT, with two native cysteines, C681^ICL2^ and C691^4.30^, labeled (Figure 3d, Extended Data Figure 10d) and mutant mGlu5 with only C681^ICL2^ labeled (Extended Data Figure 10e, f). Notably, the two bimane-labeled constructs yielded similar results (Extended Data Figure 10d, f). The addition of Quis alone did not produce a significant change (Figure 3d, Extended Data Figure 10d) in the bimane fluorescence spectrum, suggesting that an orthosteric agonist alone has a limited ability to stabilize an active conformation of TM intracellular loops. However, the addition of CDPPB to apo- or Quis-bound mGlu5 increased fluorescence, indicating a change in the ICL2 environment (Figure 3d, Extended Data Figure 10d-f). Interestingly, compared to Quis-alone, the structure of mGlu5 with Quis and CDPPB-bound, shows some changes in TM3 and TM4. These changes could be potentially contributing to changes in ICL2 (Extended Data Figure 10g). No further change was observed with the addition of G_q_ (Figure 3d). On the other hand, the mGlu5 NAM MTEP (with Quis or antagonist, LY341495) binding dramatically decreased fluorescent intensity (Figure 3d, Extended Data Figure 10d-f). The decrease in fluorescence with the NAM could be due to bimane quenching by adjacent aromatic residues^17^, such as, for example when ICL2 residue Y757^5.57^ approaches the TM domain in the active conformation. This contrasts with the increase in fluorescent intensity in the presence of the PAM, CDPPB, when ICL2 may adopt an extended conformation (ICL2 in Intermediate 1b, 2b and 3a, 3b in Figure 1a). Hence, allosteric modulators appear to regulate ICL2 conformation.

### Activation dynamics of mGlu5

Agonist binding to the VFT triggers TM rearrangements via the CRD. To investigate the mechanism of mGlu5 activation, we studied the conformational dynamics of the CRDs using single-molecule fluorescence resonance energy transfer (smFRET)^18,19^. smFRET has been used to study the VFT conformational changes upon glutamate binding^20^. We site-specifically labeled a single introduced Cys (in a minimal Cys background (Extended Data Figure 10a-c)) in the CRD of each protomer at position 560, with cysteine-reactive versions of LD555 (donor) and LD655 (acceptor), to probe the distance between the CRDs as a measure of mGlu5 activation (Figure 4a, Extended Data Figure 11a).

**Fig 4:**
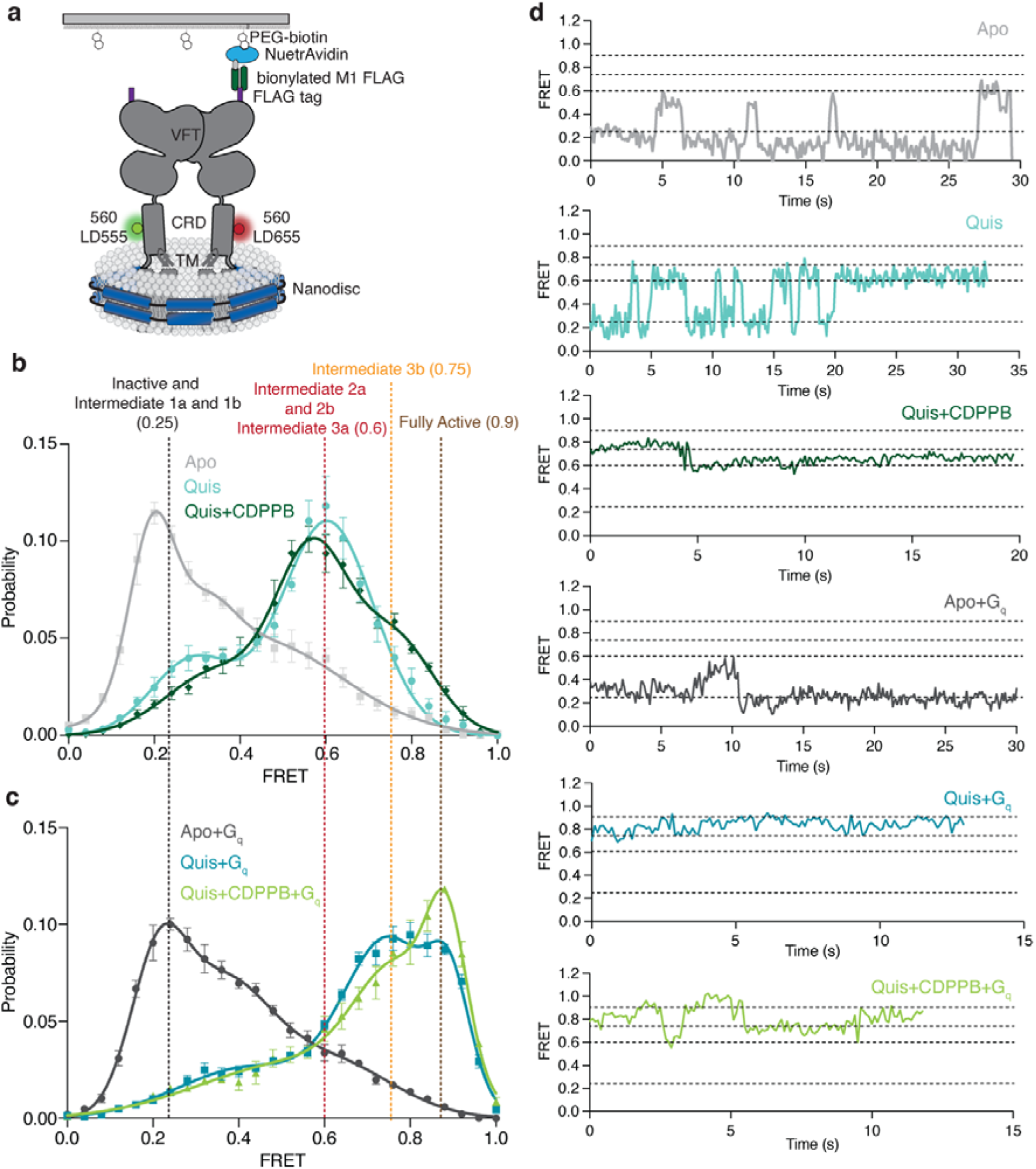
Ligand stabilised conformations of mGlu5 in nanodisc. a) A schematic representation of the smFRET experiment. b) In the Apo state (grey) a dominant inactive FRET peak at ~ 0.25 is observed (N=319). The binding of the agonist, Quis results in the appearance of a ~ 0.6 FRET state (Intermediate 2a, cyan, N=392) with a minor peak at ~ 0.25 (Intermediate 1a). The addition of CDPPB to Quis-bound mGlu5 stabilizes the ~ 0.6 FRET state (Intermediate 3a), decreases the occupancy of the ~ 0.25 state, and results in the appearance of a new FRET peak at ~ 0.75 (Intermediate 3b, dark green, N=329). High FRET (~ 0.6 and ~ 0.75) represents the active state population of the receptor with the CRDs and TMs in close proximity. Histograms are shown with a 3-Gaussian fit to the data and represented as mean ± SEM. c) The coupling of G_q_ to Apo (dark grey, N=329) remains largely unchanged compared to Apo alone (Figure 4b), while the addition of G_q_ in the presence of Quis results in the near complete abrogation of the ~ 0.6 FRET peak in favor of the ~ 0.75 peak (Intermediate 3b) and a new peak at ~ 0.9 (Fully Active) (teal, N=306), which is further stabilized in the presence of CDPPB (light green, N=317). Histograms are shown with a 3-Gaussian fit to the data and represented as mean ± SEM. d) Example FRET traces are shown for each ligand condition.

In the Apo state, mGlu5 exhibits a dominant peak centered at a FRET efficiency of ~ 0.25 and includes a broad right skew to higher FRET values (Figure 4b, d, grey, Extended Data Figure 11b). This low-FRET state at ~ 0.25 is increased by the orthosteric antagonist LY341495, indicating that it corresponds to the inactive state (Extended Data Figure 11c). Upon Quis binding, there is a shift in the occupancy to a mid-FRET state, centered at a peak of ~ 0.6 FRET (Figure 4b, d, cyan). Based on the distance between residue 560 in each protomer in the different structures that we obtained, the low and mid-FRET populations appear to correspond to the two distinct Quis-bound structures seen above (Figure 1a): the Intermediate 1a state (low-FRET) and the Intermediate 2a state (mid-FRET) (Extended Data Figure 11a). mGlu5 incubated with CDPPB alone occupies a dominant peak at ~ 0.25 (Intermediate 1b, Figure 1a) and a peak at ~ 0.6 (Intermediate 2b, Figure 1a) (Extended Data Figure 11d). Intermediate 2b was not detected by cryoEM. In the presence of Quis and CDPPB, the ~ 0.6 peak is dominant (Intermediate 3a, Figure 1a), with a decrease in the low-FRET state (Figure 4b, d dark green) and an emergence of a high-FRET peak at ~ 0.75 (Figure 4b, d dark green). This ~ 0.75 peak corresponds to a distinct active-like state of the receptor (Intermediate 3b, Figure 1a), not seen in the cryoEM structures. However, our 3Dflex analysis of the Quis and CDPPB cryoEM dataset, reveals conformations where the CRDs of the homodimers are closer (“squeeze” motion) than that seen in the Quis structure (Extended Data Figure 11e, SI Video 2), agreeing with Intermediate 3b. These observations indicate that the CRDs are dynamic, that agonist and PAM binding progressively increases the occupancy of states in which the CRDs come into closer, and closer contact, and suggests that our intermediate structures lie on this pathway. Moreover, the results show that positive allosteric modulators actuate an allosteric back communication from the TMs to the CRDs.

To study the effects of G protein on receptor conformational dynamics, we added G_q_ to Quis-bound mGlu5. The addition of G_q_ depletes the FRET peak at ~ 0.6 and shifts the receptor to inhabit the ~ 0.75 FRET state observed in Quis and CDPPB and a new FRET state at ~ 0.9 (Figure 4c, d, teal). This higher FRET peak is also seen when G_q_ is added to CDPPB-bound mGlu5 (in the absence of Quis) (Extended Data Figure 11d), agreeing with prior evidence that CDPPB is an agonist-PAM^7^. This peak at ~ 0.9 is stabilized further when G_q_ is added to mGlu5 bound to CDPPB and Quis, (Fully Active, Figure 1a) (Figure 4c, d, dark green). Though G_q_ can interact with both the Intermediate 2a (FRET ~ 0.6) and Intermediate 3a (FRET ~ 0.75) states, this ~ 0.9 high FRET, Fully Active, state is likely to be a conformation of mGlu5 stabilized only in the presence of G protein (Figure 4b, c (example traces are shown in Extended Data Figure 11e)). Though the structure of mGlu5-G_q_ has not been reported, structures of mGlu2 have been determined in the presence and absence of G protein, and no difference is observed between these mGlu2 structures^12,20^. However, our smFRET data show that, at least for mGlu5, there exists a distinct conformation in the presence of G protein.

## Discussion

A long-standing interest in the Family C GPCR field is to understand how the signal to activate is allosterically communicated over a distance of 120 Å from the orthosteric agonist binding site to the TM domain, which contains both the allosteric ligand binding pocket and the G protein binding site. We propose an activation model of mGlu5 and illustrate the effects of agonist, PAM, and G protein on the functional states that range from inactive, to fully active through several intermediates. Upon agonist binding, the upper lobes of the VFT close, while the lower lobes, CRDs, and TMs remain separated (Intermediate 1a, Figure 1a) (Figure 2a, Figure 4b). This Quis-bound Intermediate 1a conformation in mGlu5 is different from the previously observed/proposed intermediate state in other mGlus, where the agonist is bound only to one of the protomers^6,18^. The Quis-bound mGlu5 Intermediate 1a is in equilibrium with the Quis-bound Intermediate 2a (Figure 1a) conformation, with the lower lobes of the VFT, CRDs, and the TMs in close proximity^2^ (Figure 2c, Figure 4b). The addition of CDPPB results in asymmetric binding in the presence of Quis (in contrast to the symmetric binding seen in the absence of agonists) and further stabilizes an active conformation of mGlu5 (Intermediate 3a, Figure 1a) (Figure 3a, Figure 4b). Structurally, Intermediate 3a resembles the Intermediate 2a state (Figure 3b), except in the conformation of ICL2 (Figure 3d). Also, in the presence of Quis and CDPPB, there is evidence for a conformation of mGlu5 with reduced intersubunit distance, Intermediate 3b (Figure 1a, Figure 4b). The addition of G protein stabilizes a unique signaling conformation as seen from the smFRET data (Fully Active, Figure 1a)(Figure 4b). Previously, smFRET studies have been carried out on CRD-labeled mGlu2 within detergent micelles in the presence of only ligands (G protein was not used)^18,19^. These studies have shown that the activation of mGlu2 with ligands occurs by transitioning through four states and the addition of the PAM does not stabilize a new state but rather increases the occupancy of the active states. Similarly, in mGlu5, PAM alone and agonist alone do not appear to stabilize different states (Extended Data Figure 11d). However, unlike with mGlu2, PAM addition to agonist-bound mGlu5 stabilizes a state not seen with agonist alone (Intermediate 3a, Figure 1a) (Figure 4b). Further, the presence of G protein stabilizes a unique state not seen with ligands alone (Fully active, Figure 1a)(Figure 4b), which is yet to be observed structurally. This perhaps indicates slightly different activation intermediates/pathway between different mGlu receptors.

In conclusion, the combined structural and dynamic data highlight the allosteric nature of mGlu5 activation, shedding light on the conformational diversity in receptor activation and its interactions with the G protein.

## Methods

### mGlu5 purification

mGlu5 purification was carried out as previously described^2^. Briefly, human mGlu5 (21–872) with the haemagglutinin (HA) signal peptide, followed by a FLAG epitope tag (DYKDDDD) in the N terminus and a hexahistidine tag at the C terminus was expressed in sf9 cells using the Bac-to-Bac baculovirus expression system (Invitrogen). SF9 cells at density a of 3.5 X 10^6^ cells per milliliter were infected with mGlu5 virus grown for 48h at 27°C. Cells were harvested by centrifugation and lysed in a hypotonic buffer containing 10mM Tris at pH 7.8 and 1mM EDTA with protease inhibitors. After centrifugation, the pellet was solubilized with 1% (w/v) n-dodecyl-β-D-maltoside (DDM) (Anatrace), 0.1% (w/v) Cholesteryl hemisuccinate (CHS) (Steraloids), 0.2% (w/v) Sodium Cholate (Anatrace), 20 mM HEPES pH 7.5, 750mM NaCl, 30% Glycerol, Iodoacetimide 2mg/ml, protease inhibitor and 10μM MTEP for 1.5 hours at 4°C. Ca2+ was added and the supernatant after centrifugation was incubated with anti-Flag M1 affinity resin for 2 hours at 4°C. The resin was washed with 0.1% (w/v) DDM, 0.01% (w/v) CHS, 500mM NaCl, 20 HEPES pH 7.5, 2 mM Ca^2+^,10μM MTEP, followed by 0.1% (w/v) DDM, 0.01% (w/v) CHS, 100mM NaCl, HEPES pH 7.5, 2 mM Ca^2+^ and 10μM MTEP. To exchange detergent to GDN, the column was washed with an increasing concentration of GDN and a decreased concentration of DDM. Finally, the column is washed with 0.2% (w/v) GDN, 0.002%(w/v) CHS, 100mM NaCl HEPES pH 7.5, 2 mM Ca^2+^ and 10 nM MTEP. The protein was eluted in 20mM HEPES pH 7.5, 100mM NaCl, 0.2% (w/v) GDN, 0.002%(w/v) CHS, 200 μg/ml Flag peptide, 5mM EDTA, and 10 nM MTEP. The eluted protein was concentrated in a 50 kDa cut-off Vivaspin (Millipore) and run on a Superose 6 size exclusion column (GE Healthcare) in 20mM HEPES pH 7.5, 100mM NaCl, 0.2% (w/v) GDN, 0.002%(w/v) CHS and 10 nM MTEP. Fractions containing mGlu5 were concentrated, flash frozen, and stored at −80°C.

### Purification of Nb43

Nb43 was purified as described previously^2^. Nb43 in a modified pE-SUMO vector with a PelB leader sequence and SUMO fusion tag was transformed into BL21 E. coli, grown to an OD_600_ of ~ 0.6 at 37 °C and induced with 1mM IPTG. The flasks were transferred to 25 °C and allowed to express overnight (~12 hours). Bacteria were harvested and Nb43 was purified from the periplasm. Cells were thawed with SET buffer (0.5 M sucrose, 0.5 mM EDTA and 0.2M Tris pH 8.0) and stirred until homogenized. This was followed by the addition of three volumes of room-temperature milli-Q water with rapid stirring for 45 min to release the periplasmic contents. Centrifugation was performed to remove cell debris and the supernatant, after the addition of 100 mM NaCl and 10 mM MgCl_2_, was loaded onto a Ni-NTA resin. The resin was washed with 500 mM NaCl, 20 mM HEPES pH 7.5 and 20 mM imidazole, followed by 100 mM NaCl, 20 mM HEPES pH 7.5 and 20 mM imidazole. The SUMO-Nb43 was eluted in 100 mM NaCl, 20 mM HEPES pH 7.5 and 250 mM imidazole and, his-SUMO tag was removed by the addition of ULP1. Protein was dialysed overnight into 100 mM NaCl and 20 mM HEPES pH 7.5 at 4°C. Reverse Ni was performed to remove contaminants and uncleaved protein. Finally, Nb43 was subjected to size exclusion chromatography on a Superdex 200 10/30 column in 100 mM NaCl and 20 mM HEPES pH 7.5. Monomeric fractions were pooled, concentrated, and flash-frozen in liquid nitrogen.

### Nanodisc incorporation

mGlu5 in GDN was incorporated into MSP2N2 discs using the following ratio: 0.2 Receptor: 1 MSP2N2: 120 POPC: POPG (3 POPC: 2 POPG ratio). After 2 hours of incubation on ice, biobeads were added at a ratio of 1:8 mg biobeads: lipids and incubated with shaking at 4°C for 2 hours. The same amount of biobeads was added again and further incubated with shaking at 4°C overnight. The reconstitution mixture was separated from beads and applied on an M1-antiFLAG column. After washing with 100mM NaCl, 20 HEPES pH 7.5 and 15 mM Ca^2+^ to remove empty discs, the reconstituted protein was eluted in 100mM NaCl, 20 HEPES pH 7.5, 200 μg/ml Flag peptide, and 5mM EDTA. Nanodisc incorporated mGlu5 was concentrated, and injected on a Superose 6 10/30 gel filtration column in 100mM NaCl, 20mM Hepes pH 7.5. Monomeric peak fractions were collected and concentrated to ~ 5mg/mL for imaging.

### GTP turnover assay

Analysis of GTP turnover was performed by using a modified protocol of the GTPase-Glo^TM^ assay (Promega) described previously^21^. In the presence (20 uM) or absence of ligand, mGlu5 (1 μM in GDN and 0.5 μM in nanodisc) and G_i_ (1 μM for mGlu5 in GDN and 0.5 μM for mGlu5 in Nanodisc) was mixed in 20 mM HEPES, pH 7.5, 50 mM NaCl, 0.01% GDN/ 0.001% CHS (or no detergent for nanodisc sample), 100 μM TCEP, 10 μM GDP and 10 μM GTP and incubated at room temperature for 120 minutes. GTPase-Glo-reagent was added to the sample and incubated for 30 minutes. Luminescence was measured after the addition of detection reagent and incubation for 10 min at room temperature using a *SpectraMax Paradigm* plate reader.

### HDX-MS

#### Hydrogen–deuterium exchange labeling reaction

Purified mGlu5 at 60 µM [monomer] was diluted to 20 µM [monomer] in 100 mM NaCl, 20 mM HEPES, 0.05% GDN/0.005% CHS to match the concentration of mGlu5 in nanodisc (also 20 µM [monomer]). Subsequently, mGlu5 in detergent was further diluted 1:1 to 10 µM [monomer] with 10 mM monosodium glutamate, 100 mM NaCl, 20 mM HEPES, 0.05% GDN/0.005% CHS, pH 7.5, such that the final concentration of glutamate in this sample was 5 mM. An equivalent mGlu5 sample in detergent was also prepared in the absence of glutamate. Similarly, mGlu5 in nanodiscs was diluted 1:1 to 10 µM [monomer] with 10 mM monosodium glutamate, 100 mM NaCl, 20 mM HEPES, pH 7.5. All samples were incubated for 30 minutes at room temperature.

To prepare deuterated buffer, 5 mL of 10X buffer of 1 M NaCl, 200 mM HEPES, with or without 100 mM monosodium glutamate, at pH 7.5 was lyophilized overnight and then resuspended in D_2_O. 1X buffers were prepared by a 1:9 dilution with D_2_O, and a buffer of 5 mM monosodium glutamate was prepared by combining those same buffers in a 1:1 ratio. To prepare quench buffer, 27 mg of zirconium (IV) oxide, used for lipid extraction, was combined with 1 mL of quench buffer (3 M urea, 20 mM TCEP, pH 2.4), vortexed, and left on ice.

To initiate exchange, samples were diluted 1:10 into D_2_O buffer and quenched 1:1 with cold quench buffer for a total sample volume of 80 µL and left on ice. At each time point, 0.8 µL of porcine pepsin (10 mg/ml; Sigma Aldrich) and 0.8 µL of aspergillopepsin (10 mg/ml; Sigma Aldrich) were added to each sample, which was rapidly vortexed and returned to ice for 3.5 minutes. To remove zirconium (IV) oxide, each sample was transferred to a temperature-controlled centrifuge, and spun up to maximum speed (21.1 x g); the supernatant was then transferred to a new tube and flash frozen in liquid N_2_. Samples were stored at –80 °C prior to LC/MS analysis. Proteases were resuspended in 100 mM NaCl, 20 mM HEPES, pH 7.5 to 10 mg/ml and filtered (0.22 µm filter, Corning), aliquoted, flash frozen, and stored at –80 °C prior to use.

### Liquid chromatography/mass spectrometry analysis

Samples were thawed and injected into a cooled valve system (Trajan LEAP) coupled to an LC (Thermo Ultimate 3000) flowing buffer A (0.1% formic acid) at 200 µL/min. The valve chamber, trap column, and analytical column were kept at 2 °C.

Peptides were desalted for 4 minutes on a trap column (1 mM ID x 2 cm, IDEx C-128) manually packed with POROS R2 reversed-phase resin (Thermo Scientific). Peptides were then separated on a C18 analytical column (Waters Acquity UPLC BEH C18 Column, pore size 130 Å, particle size 1.7 µm, 2.1 mm ID X 50 mm) with buffer B (100% acetonitrile, 0.1% formic acid) flowing at a rate of 40 µL/min, increasing from 5% to 40% over the first 14 minutes and from 40% to 90% B over 30 s, and dropped to 5% B after 2.5 min. After 30 s, two sawtooth gradients (5% to 40% B over 30 s, 40% to 90% B over 30s, hold at 90% B for 30 s, drop to 5% B over 30 s, hold at 5% B for 30s) were performed. Peptides were eluted into a Q Exactive Orbitrap Mass Spectrometer (ThermoFisher) operating in positive ion mode (MS1 settings: resolution 140000, AGC target 3e6, maximum IT 200 ms, scan range 300-1500 m/z). For tandem mass spectrometry, mGlu5 samples were analyzed using the MS1 settings described above, with resolution 70000, and MS2 settings as follows: resolution 17500, AGC target 2e5, maximum IT 100 ms, loop count 10, isolation window 2.0 m/z, NCE 28, charge state 1 and >8 excluded, dynamic exclusion 15.0 s.

Sample time points were injected in non-consecutive order, and after every injection, a shortened, blank injection was performed to monitor for protein carryover. Briefly, blank runs were carried out with buffer A flowing at 300 µL/min. Any remaining material within the sample loop, as well as wash buffer, was desalted for 2 minutes on the same trap column; the remaining material was then separated on the same C18 analytical column flowing at a rate of 40 µL/min, increasing from 5% to 40% over the first 1.5 minutes and from 40% to 90% B over 30 s, and dropped to 5% B after 30 s. After 30 s holding at 5% B, two sawtooth gradients (5% to 40% B over 30 s, 40% to 90% B over 30s, hold at 90% B for 30 s, drop to 5% B over 30 s, hold at 5% B for 30s) were then performed.

### Peptide identification and analysis

MS2 data were processed using Byonic (Protein Metrics), resulting in a reference list of peptides, including several peptides containing glycosylation sites. Hydrogen–deuterium exchange data were analyzed in HD-Examiner (version 3.1) using default settings, and uptake values were adjusted to reflect that the final samples were 90% deuterated. Where necessary, peptide retention times histograms were manually adjusted to ensure consistency across all time points. Uptake summary data were exported from HD-Examiner and used to create uptake curves and Woods plots using Python scripts. A difference in deuteration of less than 10% between the two conditions was not considered significant, as indicated by horizontal dashed lines on Woods plots.

### Cryo-EM sample preparation and data acquisition

mGlu5 in nanodisc was incubated with Quis alone (and Nb43), Quis and CDPPB (G_q_iN and Nb43), or CDPPB alone for two hours at room temperature. For grid preparation, 3 μL of purified mGlu5 (with the different ligands) at 5 mg/ml was applied on glow-discharged holey carbon gold grids (Quantifoil R1.2/1.3, 200 mesh). The grids were blotted using a Vitrobot Mark IV (FEI) with 3 s blotting time and blot force 3 at 4°C and 100% humidity and plunge-frozen in liquid ethane.

### Image processing and 3D reconstructions

For all three samples, namely CDPPB-bound mGlu5, Quis and Nb43 bound mGlu5, and Quis+CDPPB bound mGlu5, cryo-EM data were collected on a Titan Krios electron microscope operating at 300 kV and equipped with a K3 direct electron detector. Movies were acquired at Counting mode with a calibrated pixel size of 1.111 Å/pixel and a total dose of ~51.6 electrons/Å^2^, fractionated across 50 frames (Extended Data Figure 2, 6, and 7, Extended Data Table 1).

Single particle data processing was performed using *cryoSPARC 3.3.2*^22^. Initially, motion correction and CTF estimation were carried out using *Patch Motion Corr* and *Patch CTF*, followed by template-based particle picking utilizing previously determined mGlu5 structures (EMD-0345 or EMD-0346). The detailed data processing workflows can be found in Extended Data Figures 2, 6, and 7. Picked particles were sorted using 2D classification, ab initio, and heterogenous refinement. For each structure/conformation, we applied both global non-uniform refinement^23^ and local refinement on the CRD-TM region. The final maps, generated with *UCSFChimera*^24^, are composite maps of the global non-uniform refinement and the locally refined CRD-TM region. Map sharpening was performed with *Autosharpen* within *Phenix*^25,26^.

Lastly, local resolution estimation and 3DFSC were employed to assess the local resolution and orientation distribution of the final dataset^27^.

### Model building and refinement

The initial template was mGlu5 from PDB codes 6N51 and 6N52. Ligand coordinates and geometry restraints were generated using *phenix.elbow*^25^. *Coot*^28^ was used for iterative model building and the final model was subjected to global refinement and minimization in real space using *phenix.real_space_refine* in *Phenix*^25^. FSC curves were calculated between the resulting model and the half map used for refinement as well as between the resulting model and the other half map for cross-validation (Extended Data Figures 2, 6, and 7). The final refinement parameters are provided in Extended Data Table 1.

### 3DVA and 3DFlex Analysis

*CryoSPARC’s 3D Variability Analysis* (3DVA) was used to investigate the conformational heterogeneity in the final data sets for Quis-bound or Quis + CDPPB-bound active state^14^. The particles used for final non-uniform refinement were processed by 3DVA with three modes, and a mask encompassing the whole receptor, including the nanodisc. Following 3DVA, the three principal components were subjected to 3DVA Display using simple output mode and 20 frames. The resolution was low pass filtered to 5Å to avoid noise dominating the determination of eigenvectors. The results were visualized using *UCSFChimera* (SI Video 1).

*CryoSPARC’s 3D Flexible Refinement* (3DFlex) was also used to investigate and confirm the conformational heterogeneity in the Quis+CDPPB bound active state^15^. For 3DFlex on the whole receptor, the particles used for the final non-uniform refinement were cropped into 256 pixels (1.48Å/pixel) for reconstruction and 128 pixels (2.95Å/pixel) for training. A mask, which encompassed the whole receptor including the nanodisc, was divided into 20 tetrahedral cells to prepare the mesh. For 3DFlex training, we optimized different parameters and ended up using 3 latent dimensions, 0.1 rigidity prior strength, and 10 latent centering strength. The latent distributions along 3 dimensions are visualized using *UCSFChimera* (SI Video 2).

For 3DFlex on the CRD+TM regions, the particles used for TM local refinement were kept in their original box size for data prep and cropped into 170 pixels (2.22Å/pixel) for training. A mask encompassing the TM regions (Extended Data Figure 6a), excluding the nanodisc, was divided into 40 tetrahedral cells to prepare the mesh. For 3DFlex training, we optimized different parameters and ended up using 3 latent dimensions, 0.5 rigidity prior strength, and 10 latent centering strengths. The latent distributions along 3 dimensions are visualized using *UCSFChimera* (SI Video 2).

### Minimal Cysteine expression and purification

We developed a minimal cysteine (minCys) construct of mGlu5, to enable site-specifically labeling. In mGlu5, two Cys residues, one in the intracellular end of TM4 (C691^4.30^) and the other in ICL2 (C681^ICL2^) appear to be exposed to being labeled, mutation of these to Ala largely abolished background labeling. Human mGlu5 (21–872) with the haemagglutinin (HA) signal peptide, followed by a FLAG epitope tag (DYKDDDD) in the N terminus and a hexahistidine tag at the C terminus was cloned into pcDNA-Zeo-tetO. Two native cysteine residues were mutated to make the minimal cysteine construct (C691^4.30^A and C681^ICL2^A) and for bimane spectroscopy studies a single C691^4.30^A mutant was cloned. For smFRET studies, an engineered Cys was introduced into the CRD (560C). The bimane and smFRET constructs were transfected into Expi293F (Thermo Fisher) cells stably expressing the tetracycline repressor using an Expifectamine transfection kit (Thermo Fisher) following the manufacturer’s recommendations with the following modifications. Two days post-transfection, mGlu5 expression was induced with doxycycline (4 μ/ml and 5 mM sodium butyrate) in the presence of 1 μM MTEP. Cells were harvested 30 hours post-induction and stored at −80°C until use. Further purification and nanodisc incorporation were performed following the protocol described earlier for pellets from insect cells.

### Bimane Spectroscopy

WT mGlu5 or C691^4.30^A mutant in nanodisc at 10 μM was incubated with a 10-molar excess of bimane at room temperature for one hour. The excess label was removed using size exclusion chromatography on a Superose 6 10/300 Increase column in 20 mM HEPES pH 7.5 and 100 mM NaCl. Bimane-labeled mGlu5 at 0.1 μM was incubated with ligands (10 uM) for one hour at room temperature. Fluorescence data were collected at room temperature in a 150 μL cuvette with *FluorEssence v3.8 software* on a Fluorolog instrument (*Horiba*) in photon-counting mode. Bimane fluorescence was measured by excitation at 370 nm with excitation and emission bandwidth passes of 4 nm. The emission spectra were recorded from 410 to 510 nm with a 1 nm increment and 0.1 s integration time.

### smFRET sample preparation and data collection

Nanodisc incorporated minCys mGlu5 (with residues C691^4.30^ and C681^ICL2^ mutated) with an introduced Cys at position 560 was incubated with 5-molar excess of donor (LD555) and 10-molar excess acceptor (LD655) at room temperature for 20 min. Following the incubation with 5 mM cysteine for 10 min, the sample was subjected to size exclusion chromatography on a Superose 6 10/300 Increase column in 20 mM HEPES pH 7.5 and 100 mM NaCl, for the removal of excess labels.

mPEG (Lysan Bio) was used to passivate glass coverslips (VWR) and, further doped with biotin PEG16 to inhibit nonspecific protein adsorption. The coverslips, before each experiment, were incubated with *NeutrAvidin* (Thermo Fisher), followed by 10 nM biotinylated antibody (mouse anti-FLAG, GenScript). Chambers were flushed to remove reagents, between each conjugation step. The anti-FLAG antibody was diluted and washed in 50 mM NaCl, 10 mM Tris, pH 7.5. Labeled mGlu5 was diluted ~ 100-fold and applied to coverslips to achieve optimum surface immobilization (~400 molecules in a 2,000 μm^2^ imaging area). Unbound receptors were washed away with buffer. smFRET imaging was performed in an imaging buffer consisting of 3 mM Trolox, 100 mM NaCl, 2 mM CaCl2, 20 mM HEPES pH 7.5, and an oxygen scavenging system (0.8% dextrose, 0.8 mg/ml glucose oxidase, and 0.02 mg/ml catalase). Samples were imaged with a 1.49 NA 60X objective (Olympus) on a total internal reflection fluorescence microscope with 100 ms time resolution unless stated otherwise. Lasers at 532 nm (Cobolt) and 633 nm (Melles Griot) were used for donor and acceptor excitation, respectively. Fluorescence was passed through a Chroma ET550lp filter and split into donor and acceptor signals with a Chroma T635lpxr. FRET efficiency was calculated as (IA-0.1ID)/(ID+IA), in which ID and IA are the donor and acceptor intensity, respectively, after background subtraction. Imaging was with 100 millisecond acquisition time (10 Hz) with a Photometrics Prime 95B CMOS camera.

### smFRET data processing

The fluorescence movies were analyzed with SPARTAN version 3.7^29^. Donor and acceptor channels were aligned using the first 10 frames of each movie while excluding particles closer than 3.5 pixels using an integration window of 12 pixels. Traces showing single-donor and single-acceptor photobleaching with a stable total intensity for longer than 5 seconds (50 frames), SNRbg > 15, and donor/acceptor correlation coefficients < 0.0 were collected (20–30% of total molecules per imaging area). Nonlinear filter^30^ was used for smoothing individual traces with the following filter parameters: window = 2, M = 2, and P = 15 for histograms. smFRET histograms were compiled from ≥100 molecules per condition (100 millisecond time resolution). Error bars in the histograms represent SEM from ≥4 independent movies. To ensure that traces of different lengths contribute equally, histograms from individual traces were normalized to one before compiling. Histograms were fit to 3 Gaussians (based on the Akaike information criteria, Figure 11b) using *Graphpad Prism 9*.

## Data availability

The data that support this study are available from the corresponding authors upon request. The cryo-EM density maps have been deposited in the Electron Microscopy Data Bank (EMDB) under accession codes EMDB-41092, EMDB-41099, EMDB-41139 and EMDB-41069. Model coordinates have been deposited in the Protein Data Bank (PDB) under accession number 8T7H, 8T8M, 8TAO, 8T6J.

## Acknowledgements

This work was supported by the National Institutes of Health grant R01NS119826 (E.Y.I.). We would like to thank National Institute of Drug Abuse (NIDA). E.Y.I. is a Weill Neurohub Investigator. B.K.K. is a Chan Zuckerberg Biohub Investigator.

## Author Contributions

KKK and BKK conceived the project. KKK prepared samples, froze grids and collected cryoEM data with help from CZ and EM. KKK and JX developed the minimal cysteine mGlu5 construct and performed bimane studies and, made smFRET samples with help from ESO.

HW processed cryoEM data and performed the 3DVA and 3dflex analysis.

CH collected and analysed smFRET data under the supervision of EYI.

NRL performed and analysed the HX data under the supervision of SM.

AK helped with structure analysis.

KKK, HW and BKK wrote the manuscript with inputs from all the authors.

## Extended Data Figures

**Extended Data Figure 1:**
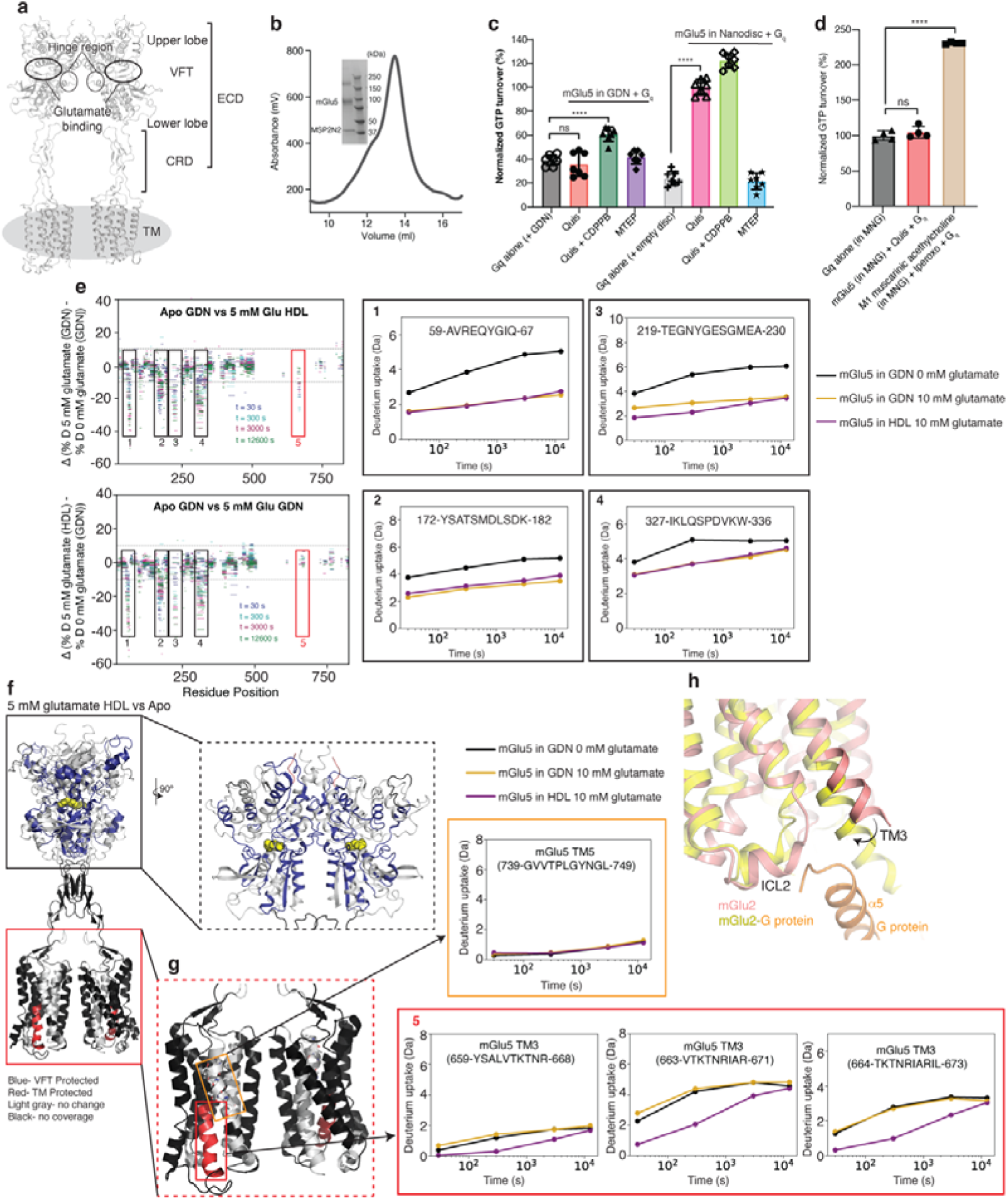
mGlu5 activation in detergent compared to lipid environment. (a) Structural domains of mGlu5. (b) SEC trace and SDS-PAGE gel of mGlu5 in nanodisc. (c) GTP turnover assay showing mGlu5 induced G_q_ turnover. In the presence of agonist Quis, mGlu5 in detergent does not induce significant G_q_ (red) turnover compared to G_q_ alone (grey). The addition of Quis and CDPPB (dark green) to mGlu5 in detergent results in a small but significant increase in G protein turnover. With mGlu5 in nanodiscs, the addition of Quis significantly increases G_q_ turnover (magenta). Quis and CDPPB (light green) further increase the GTP turnover of G_q_. The negative allosteric modulator, MTEP inhibits turnover in mGlu5 nanodiscs condition (blue). Data represented as mean ± SD, ns= 0.4124, *p* < 0.0001****, unpaired *t*-test (two-tailed), n= 7 individual experiments (data normalization was done with the average value of Quis-bound mGlu5 in nanodiscs as 100% and receptor alone as 0%). (d) In the presence of the agonist iperoxo, muscarinic acetylcholine M1 receptor (in MNG) induces significant GTP turnover in G_q_ (*p* < 0.0001****). But no difference is seen with Quis-bound mGlu5 (in MNG) and G_q_ (ns= 0.5374). Data represented as mean ± SD, *p* values are from unpaired *t*-test (two-tailed), n= 4 individual experiments. Data normalization was done with the average value of G_q_ in MNG as 100% and buffer alone as 0%. (e) HDX-MS data is plotted as the difference in the percent deuteration for a given peptide at a given time point against the sequence position for Apo mGlu5 in detergent (GDN) vs 5 mM glutamate-bound mGlu5 in detergent (GDN) (top) and Apo mGlu5 in detergent (GDN) vs 5 mM glutamate-bound mGlu5 in nanodisc (HDL) (bottom). Black boxes numbered 1-4 are example regions in the VFT that show no difference between mGlu5 in detergent (GDN) and nanodisc (HDL) (the corresponding deuterium uptake plots are shown on the right). The red box is a region in the TM that shows a difference between agonist-bound mGlu5 in detergent and nanodisc (HDX-MS exchange curves shown in Extended Figure 1g). (f) HDX-MS changes in Apo mGlu5 in detergent and agonist-bound mGlu5 in nanodisc are plotted onto the mGlu5 structure (PDB code: 6N51). (g) Colored red is the TM3 region, where peptides were observed in HDX-MS measurements. Deuterium uptake plots of these TM3 peptides (red box in Extended Fig 1e) show that Apo (black) and receptor in GDN in the presence of 5 mM glutamate (yellow) overlay well. Whereas, TM3 peptides of mGlu5 in nanodisc (magenta) do not overlay with Apo. Shown in the orange box is a TM5 peptide showing no change in deuterium uptake between the conditions. (h) Agonist-bound mGlu2 (PDB code: 7MTR) is overlayed with agonist-bound mGlu2-G protein complex (PDB code: 7MTS) showing conformational changes in the intracellular region of TM3.

**Extended Data Figure 2:**
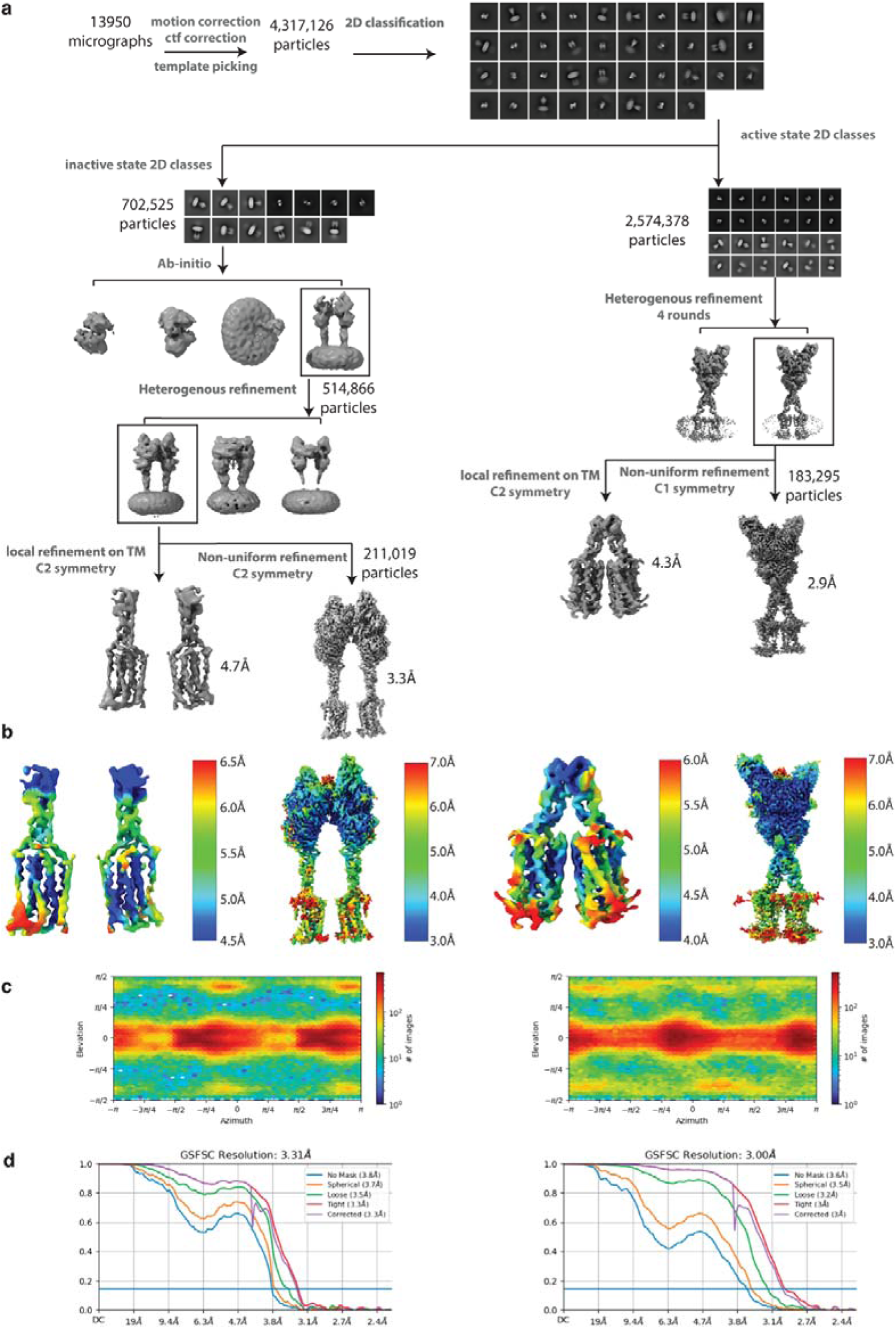
Cryo-EM data processing workflow and resolution assessment of Quis-bound maps. (a) Workflow of cryo-EM data processing to obtain Quis-bound Intermediate 1a and Quis-bound Intermediate 2a structures. (b) Local resolution maps of the Quis-bound structures. (c) Angular particle distribution of the Quis-bound structures. (d) Gold-standard FSC curves of the structures.

**Extended Data Figure 3:**
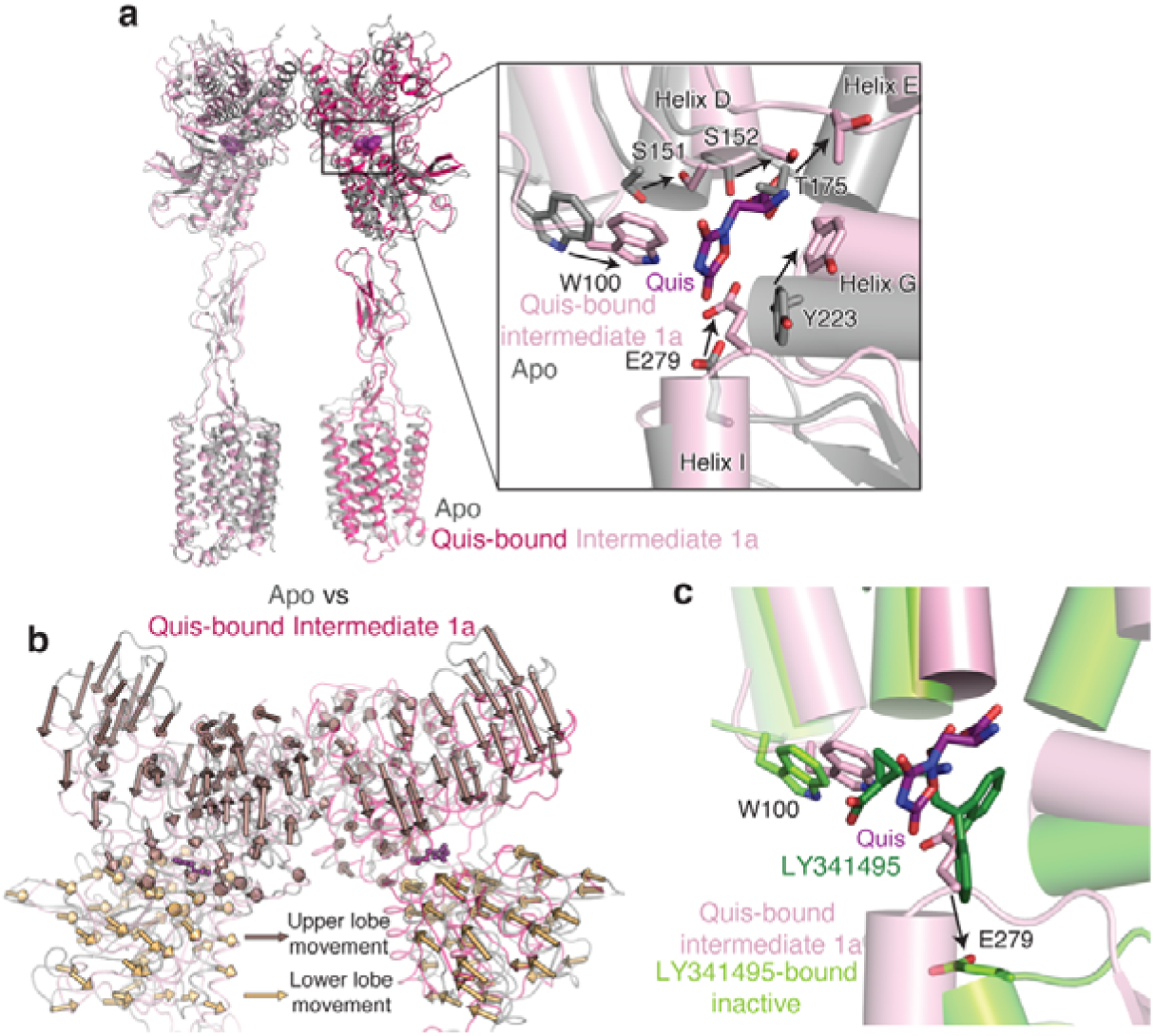
Comparison of Apo and Quis-bound Intermediate 1a structures. (a) Overlay of the Apo (grey, PDB: 6N52) and Quis-bound Intermediate 1a showing CRDs and TMs in an “inactive” state. Insert shows the Quis binding pocket. (b) Movement of the VFTs upon agonist binding in Quis-bound Intermediate 1a state compared to the Apo state (PDB 6N52, grey). Arrows represent the movement of every 5 Cα atoms from the Apo to the Intermediate 1a state upon Quis binding. Nb43 is shown in yellow. (c) To get insights into structural changes needed to initiate activation, we compared the Quis-bound Intermediate 1a (light pink) and the antagonist, LY341495-bound (PDB: 7FD9, dark green) mGlu5 structures. LY341495 binding to the receptor inhibits the movement of residues W100 and E279.

**Extended Data Figure 4:**
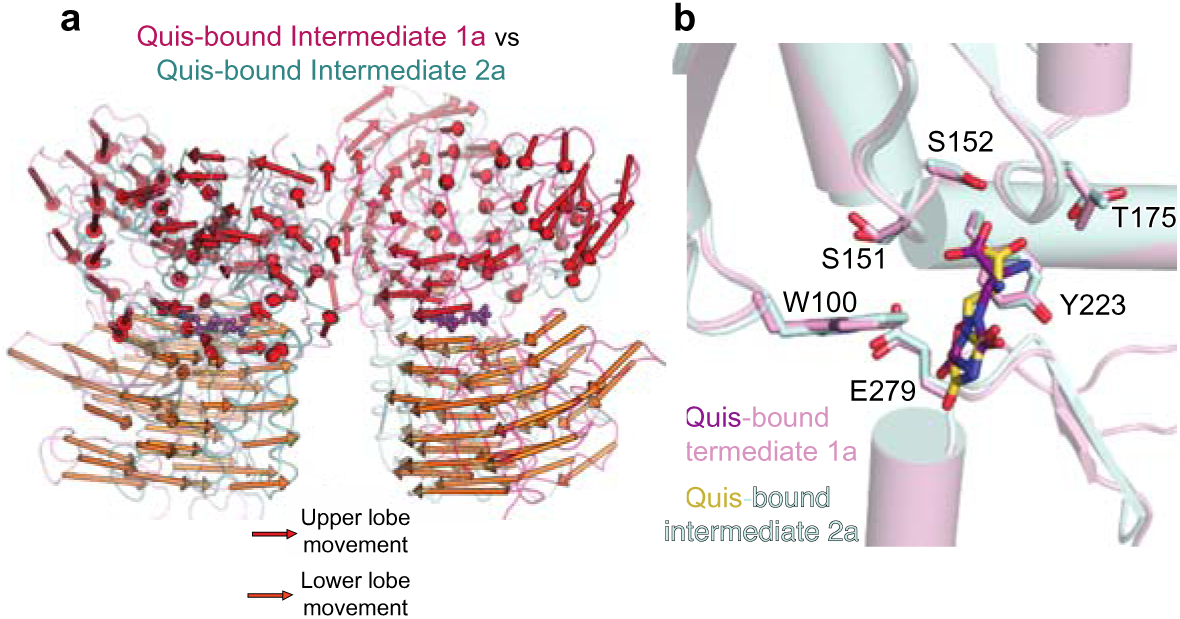
Comparison of Quis-bound Intermediate 1a and Quis-bound Intermediate 2a structures. (a) Comparing the movement of the VFTs in the Quis-bound Intermediate 1a (light pink, magenta) and the Quis-bound Intermediate 2a states (cyan and teal) show large rearrangements in the lower lobe, with relatively smaller changes in the upper lobe. Arrows represent the movement of every 5 Cα atoms from the Intermediate 1a state to the Intermediate 2a state. (b) Single protomer alignment of Quis-bound Intermediate 2a (cyan) and Quis-bound Intermediate 1a (light pink) structures show no change in the Quis binding pocket.

**Extended Data Figure 5:**
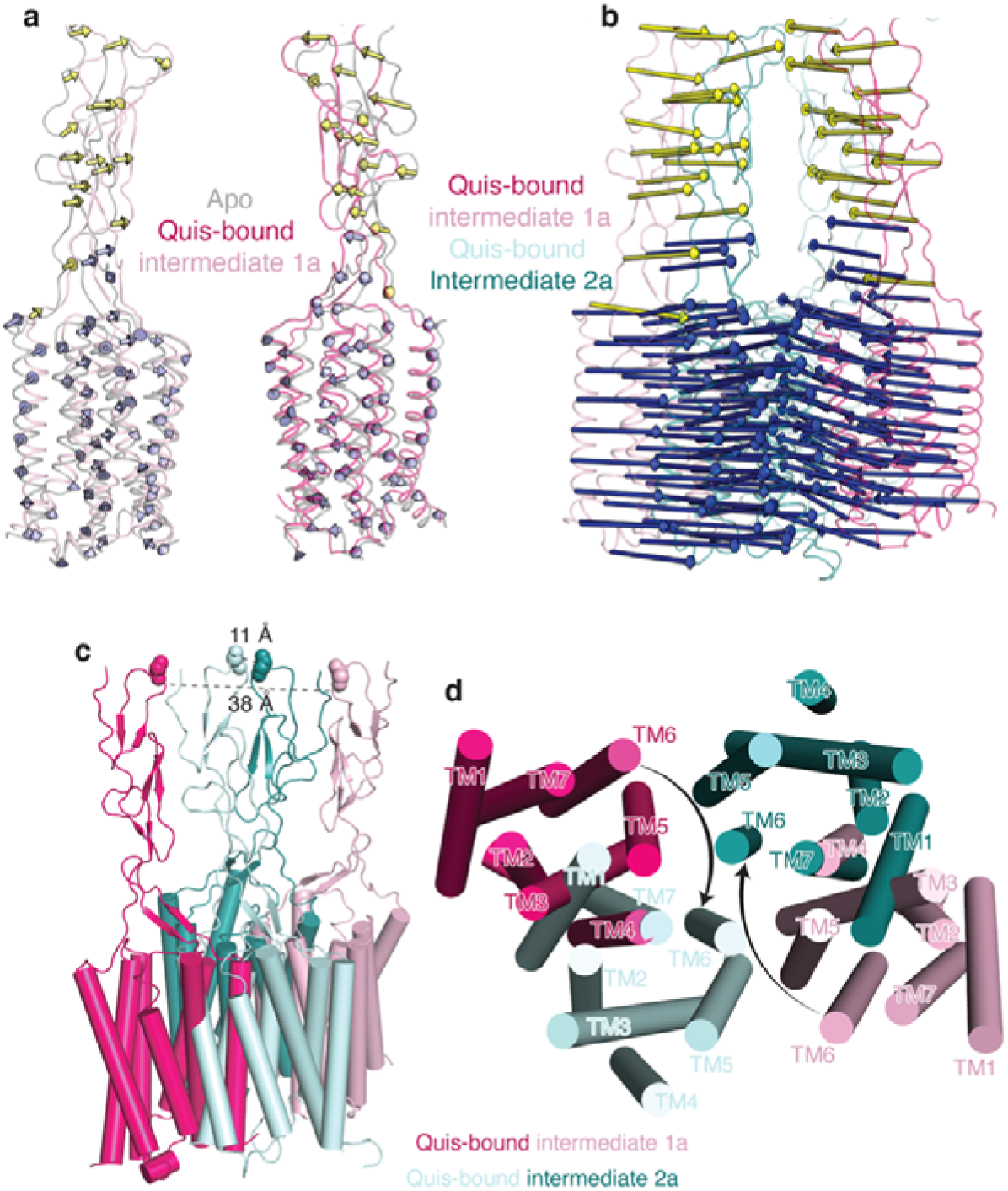
mGlu5 transmembrane changes upon activation. a) Overlay of Apo (grey, PDB: 6N52) and Quis-bound Intermediate 1a states show minimal changes in the CRDs and TMs. Arrows represent the movement of every 5 Cα atoms from Apo to Intermediate 1a. b) Large changes in the CRDs and TMs are seen when comparing the Quis-bound Intermediate 1a and the Quis-bound Intermediate 2a states. Arrows represent the movement of every 5 Cα atoms from Intermediate 1a state to Intermediate 2a. c) The CRDs in the Quis-bound Intermediate 1a structure are separated by ~ 38 Å (as measured at residue E527). In the Quis-bound Intermediate 2a state, the twisting of the lower lobe enables the CRDs (~ 11 Å at residue E527) and TMs to move adjacent to each other. d) The TMs in the Quis-bound Intermediate 1a structure are far apart with TM5 being the most proximal helix pair (~ 21 Å). In the Quis-bound Intermediate 2a state the TMs of the protomers, in addition to moving closer to each other, rotate ~ 20° to form a TM6-TM6 interface, a hallmark of Family C activation.

**Extended Data Figure 6:**
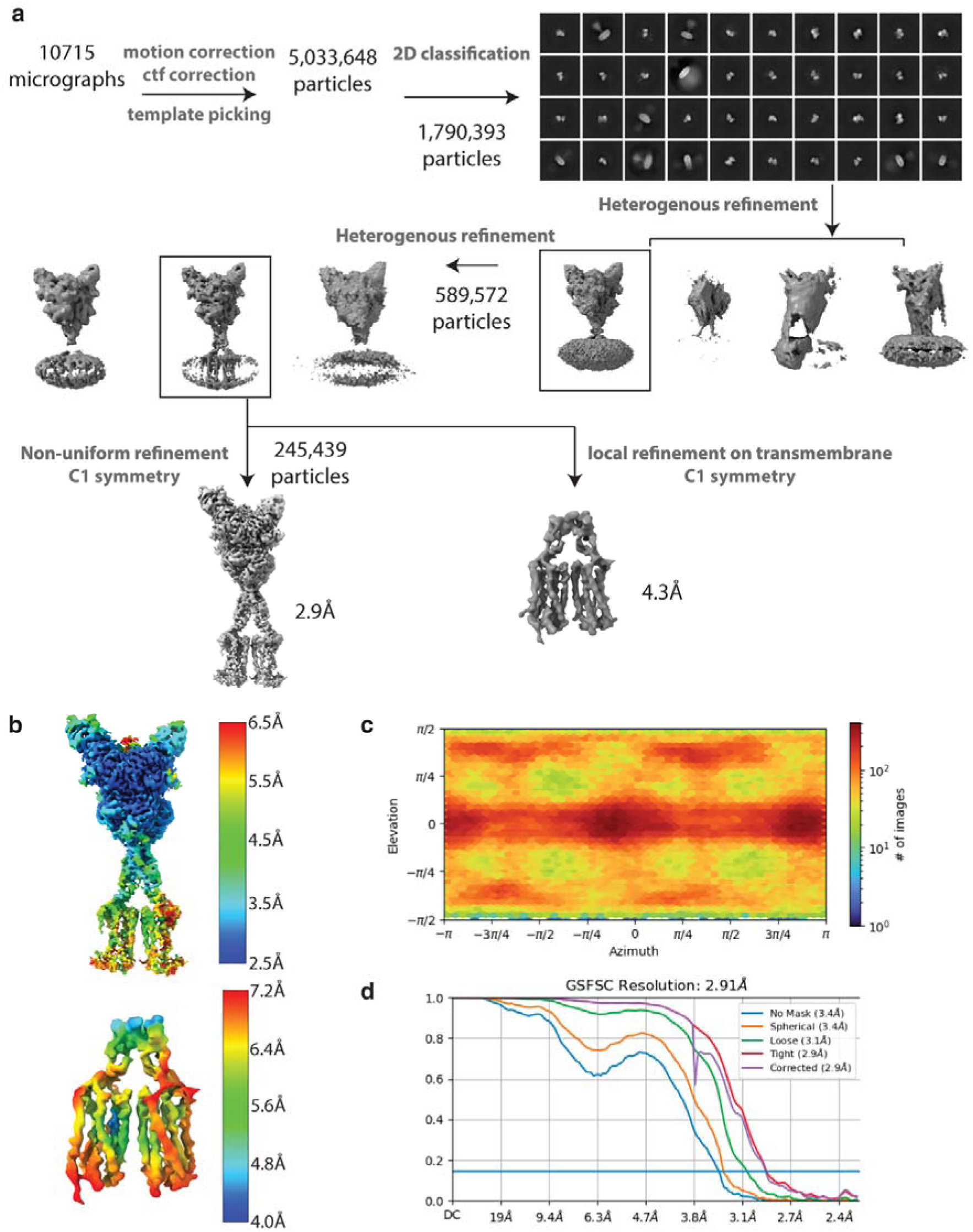
Cryo-EM data processing workflow and resolution assessment of CDPPB, Quis-bound map. (a) Workflow of cryo-EM data processing to obtain CDPPB, Quis-bound mGlu5 structure, Intermediate 3a. (b) Local resolution maps of the CDPPB, Quis-bound mGlu5 structure. (c) Angular particle distribution of the structure. (d) Gold-standard FSC curves of the Quis-bound mGlu5 structure.

**Extended Data Figure 7:**
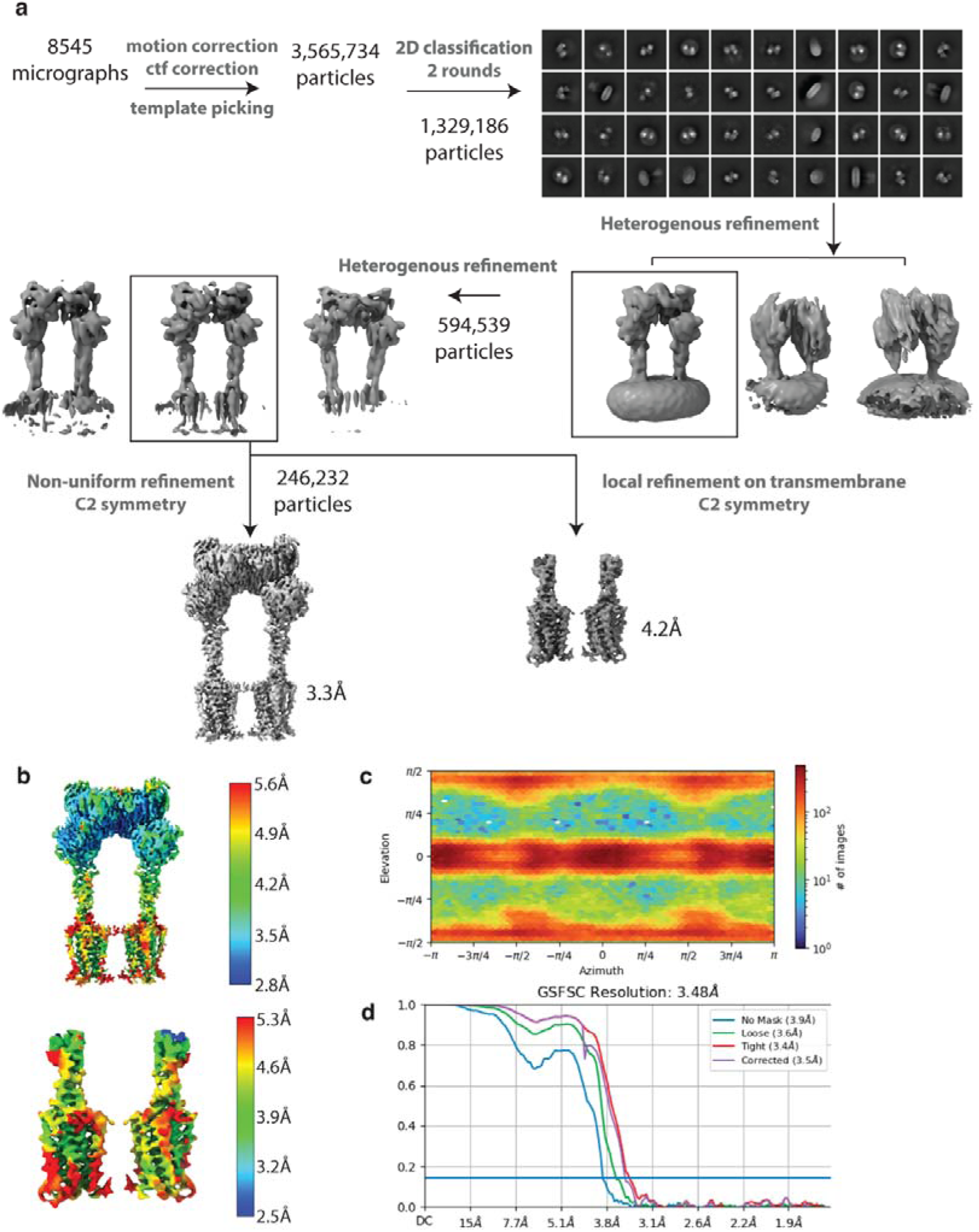
Cryo-EM data processing workflow and resolution assessment of CDPPB-bound map. (a) Workflow of cryo-EM data processing to obtain CDPPB-bound mGlu5 Intermediate 1b structure. (b) Local resolution maps of the CDPPB-bound mGlu5 structure. (c) Angular particle distribution of the structure. (d) Gold-standard FSC curves of the CDPPB-bound mGlu5 structure.

**Extended Data Figure 8:**
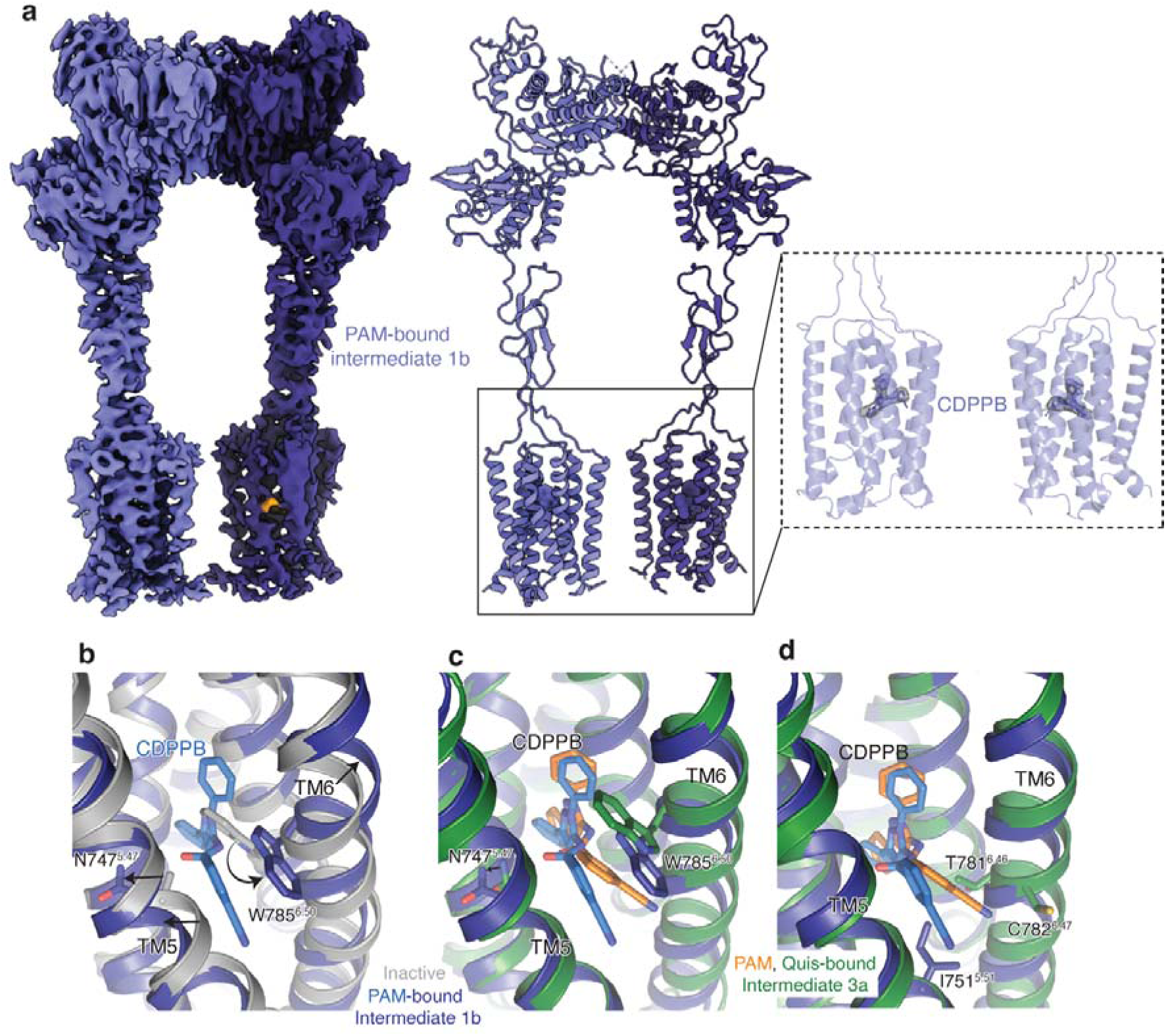
CDPPB-bound structure analysis. (a) Cryo-EM density and model of CDPPB-bound mGlu5 Intermediate 1b in a nanodisc. Also shown is the density for the two bound CDPPB, one in each TM domain. (b) Comparison of the allosteric binding pocket in Apo (PDB:6N52, grey) and CDPPB-bound mGlu5 (dark blue), shows changes in TM5 (N747^5.47^) and TM6 (W785^6.50^) to accommodate CDPPB (slate). (c) Overlay of CDPPB from Intermediate 1b (dark blue) and Intermediate 3a structures showing minimal changes in the conformation of TM5 and TM6. (d) Residues that interact with CDPPB only in Intermediate 3a are shown in green (T781^6.46^ and C782^6.47^) and those that interact with CDPPB only in Intermediate 1b are shown in blue (I751^5.51^).

**Extended Data Figure 9:**
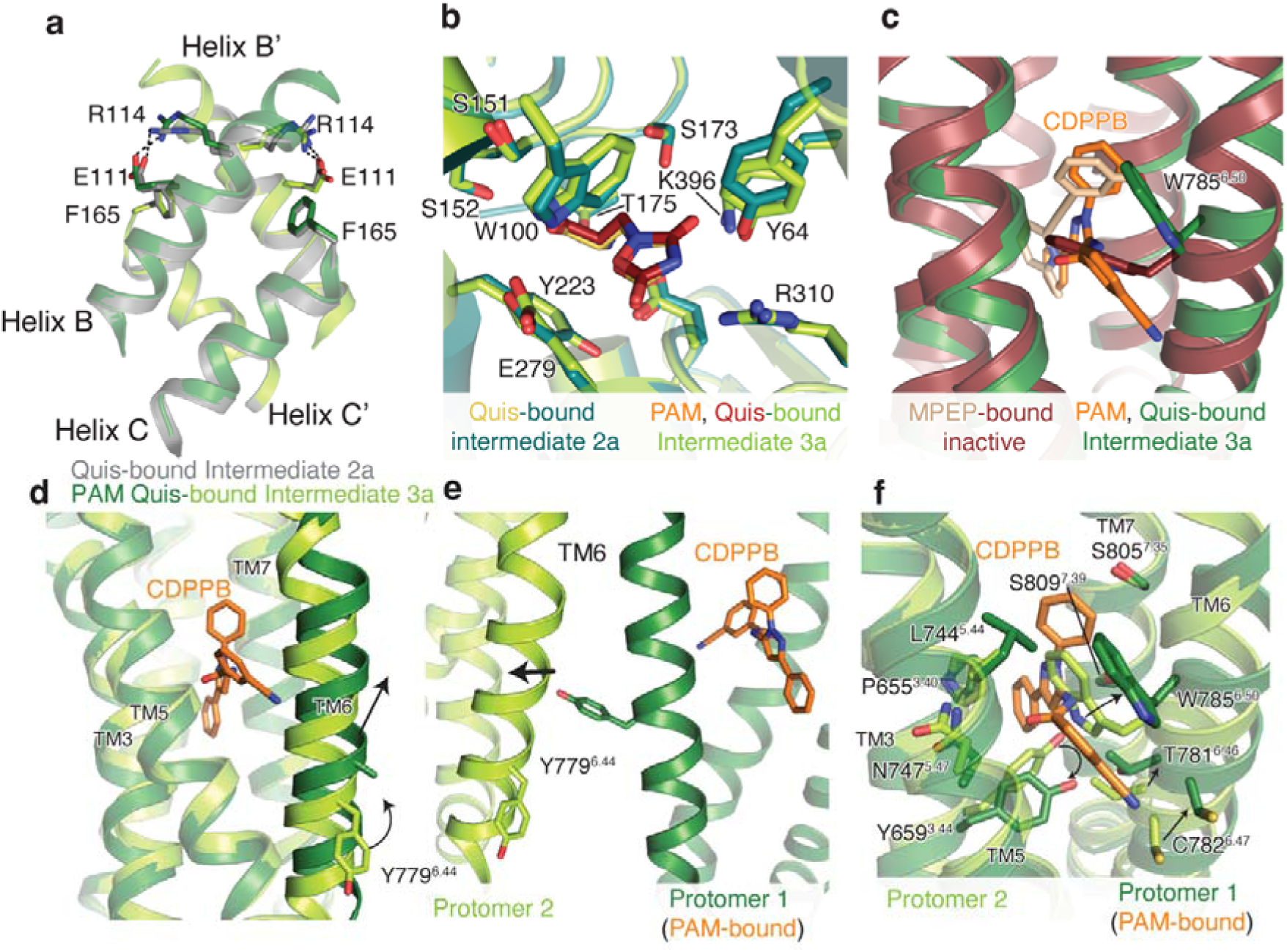
CDPPB, Quis-bound structural analysis. (a) Overlay of intersubunit B and C helices in Quis-bound Intermediate 2a state and CDPPB, Quis-bound Intermediate 3a structure. Residues R114 and E111 interact in both structures. (b) Overlay of Quis binding pocket in Quis-bound Intermediate 2a and CDPPB, Quis-bound Intermediate 3a structures, showing no difference in the ligand pocket. (c) The conformation of residue W785^6.50^ is different in the structure with the NAM, MPEP (PDB: 6FFI, brown) compared to that with the PAM, CDPPB (dark green). (d) TM6 in the CDPPB-bound Protomer 1 has moved outward compared to Protomer 2 with no CDPPB bound. In CDPPB-bound Protomer 1, Y779^6.44^ points towards the intersubunit interface, as seen in (e). Though we cannot model the Y779^6.44^ sidechain in Protomer 1 with confidence due to a lack of good density, we have added the most frequently occurring rotomer of Tyr. (f) Comparison of the allosteric pocket in CDPPB-bound protomer (protomer 1, dark green and CDPPB shown as orange) and the protomer with no CDPPB (protomer 2, green).

**Extended Data Figure 10:**
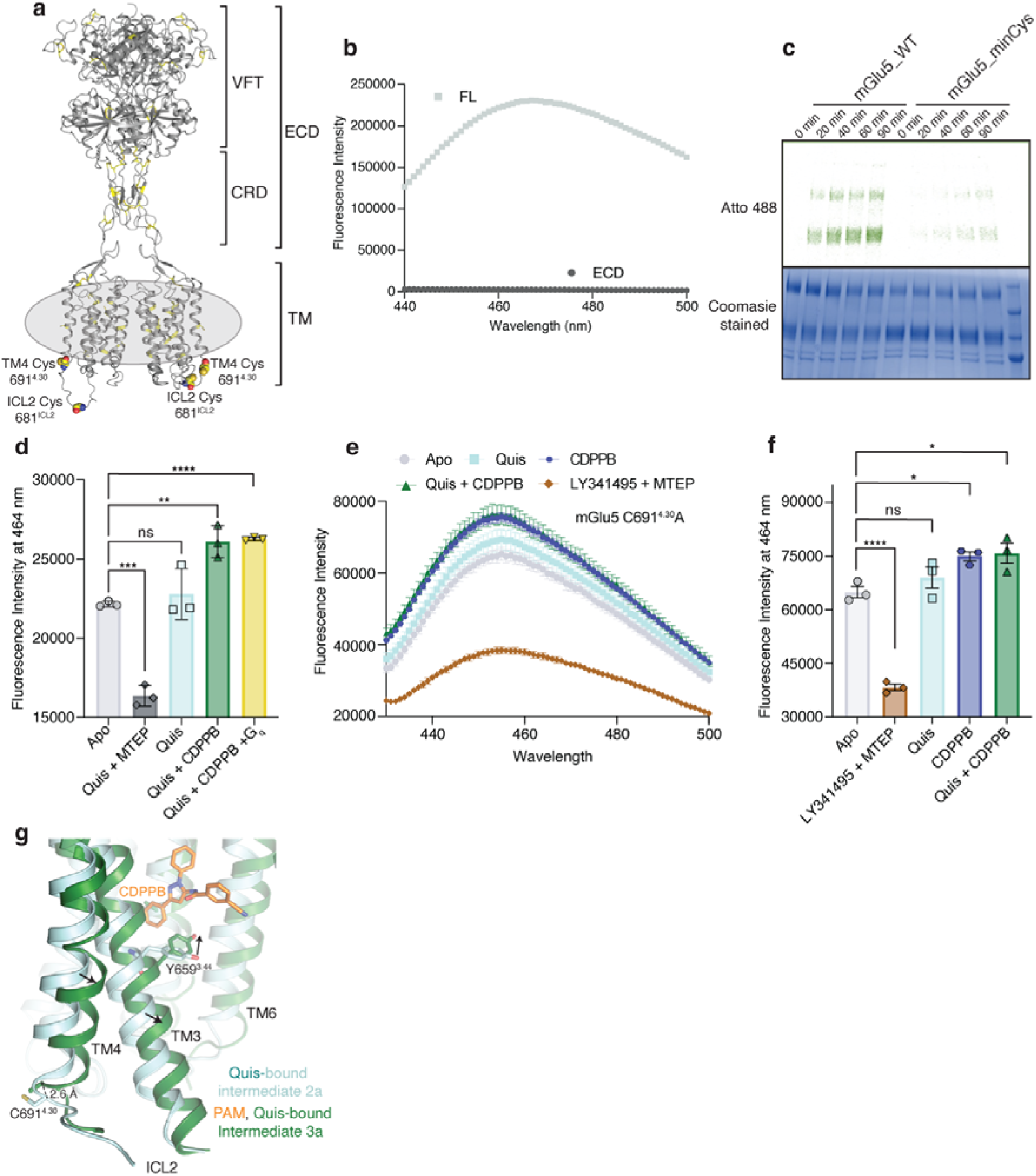
Characterisation of minimal cysteine mGlu5 and ICL2 conformation. (a) Residues Cys691^4.30^ and Cys681^ICL2^ that contribute to background labeling with dyes are shown as spheres. Other cysteine residues in the receptor are shown as yellow sticks. (b) mGlu5 full-length and ECD alone (VFT and CRD) were labeled with the cysteine reactive dye, monobromobimane. Though no signal was seen for ECD (dark grey), full-length (FL) mGlu5 produced a bimane spectrum (light grey). This implies that mGlu5 TMs have cysteine residues that are exposed to being labeled with bimane. n = 1 individual experiment. (c) WT and minimal cysteine (C691^4.30^A and C681^ICL2^A) constructs were labeled with Atto488. Unlike WT, the minimal cysteine construct exhibits almost no background labeling for the times tested. (d) Fluorescence intensity at 464 nm for mGlu5 WT labeled with bimane (reading out on ICL2 conformation from Fig 3e) is plotted for the different ligand conditions. Though there is no significant difference between Apo (light grey) and Quis (cyan), the addition of Quis and CDPPB (dark green) showed a significant change. No further change was detected with the addition of G_q_ to the Quis and CDPPB condition (yellow). The addition of MTEP resulted in a significant decrease in fluorescence intensity (dark grey). Data represented as mean ± SD, ns = 0.5326, *p=* 0.0001***, *p* = 0.0026**, *p* < 0.0001****, unpaired *t*-test (two-tailed), n = 3 individual experiments. (e) Bimane spectra of mGlu5 in nanodiscs labeled only at C681^ICL2^ (C691^4.30^A construct). Unlike adding Quis (cyan) which resulted in no change in the spectrum, the addition of CDPPB alone (blue) or Quis and CDPPB (dark green) increases the fluorescence. On the other hand, LY341495 and MTEP (brown) cause a decrease in fluorescence. Data represented as mean ± SEM, n = 3 individual experiments. (f) Plotting the fluorescence intensity at 464 nm for bimane data in Extended Data Figure 10e shows a significant difference between CDPPB alone (blue), Quis and CDPPB (dark green), and LY341495 and MTEP (brown) compared to Apo (grey). Data represented as mean ± SEM, ns = 0.5713, *p* < 0.0001, *p* = 0.0257* (Apo vs CDPPB), *p* < 0.0160* (Apo vs Quis + CDPPB), unpaired *t*-test (two-tailed), n = 3 individual experiments. (g) Comparison of Quis-bound (cyan) and CDPPB, Quis-bound structures (dark green) showing changes in TM3 and TM4. Also shown is the position of residue C691^4.30^ which is bimane labeled in the WT construct (Figure 3e, Extended Figure 10e).

**Extended Data Figure 11:**
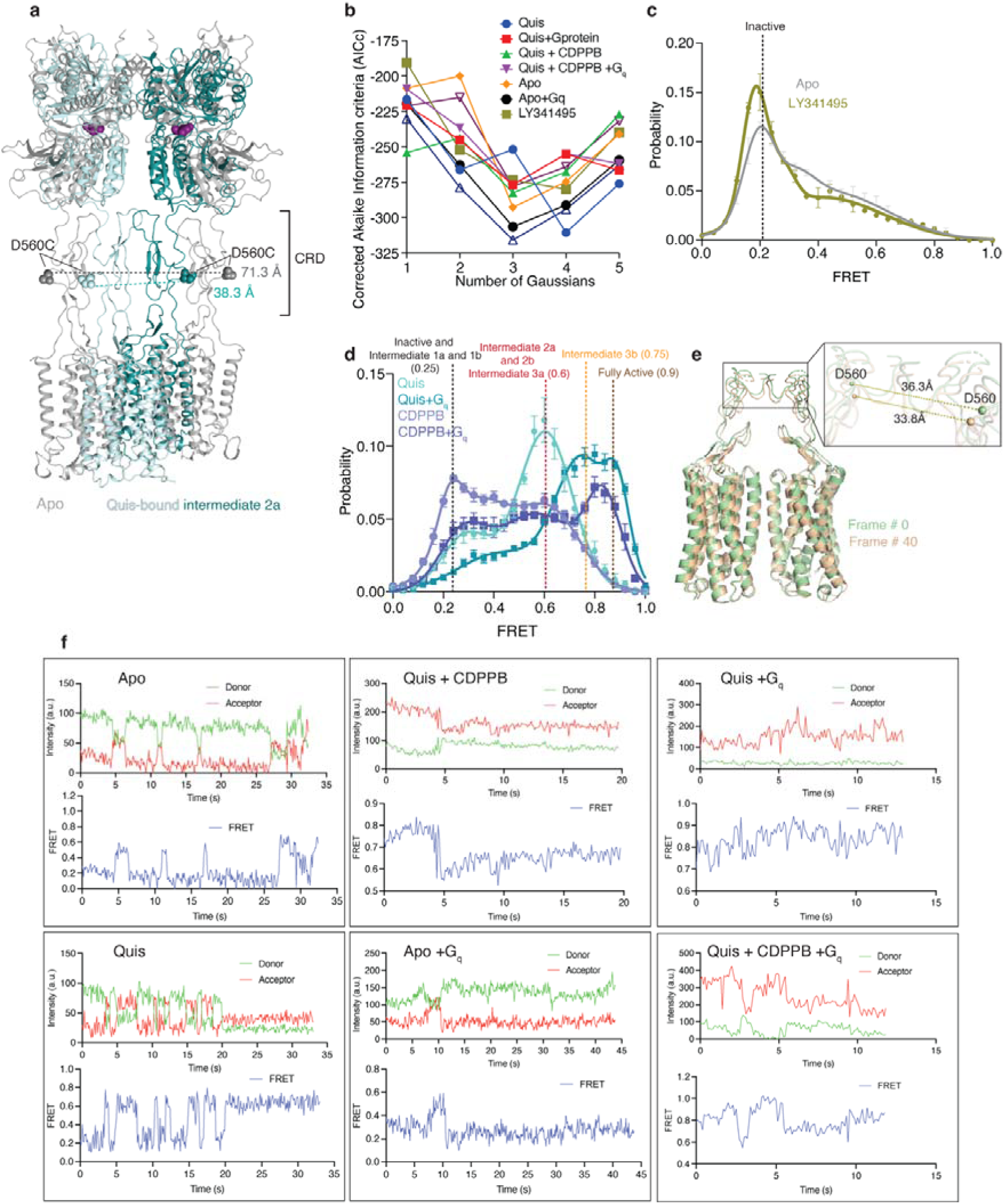
smFRET fitting statistics and analysis. (a) Interdye distance between residue D560 in Apo (grey, 71.3 Å) and Quis-bound Intermediate 2a (36.3 Å). Both these distances correlated well with the observed FRET values (Figure 4b). (b) Plot of the Akaike information criterion (corrected for small sample size, AICc) values for analysis with 1 to 5 Gaussians fits for the smFRET data. The AICc values showed broad minima at 3 and 4 fits. 3 Gaussians were used to fit the data. (c) smFRET data showing the comparison of Apo (grey, N=319) and antagonist-bound mGlu5 (brown, N=245). (d) The addition of CDPPB alone results in two FRET peaks, one at ~ 0.25, Intermediate 1b state, and the other at ~ 0.6, the Intermediate 2b state (slate, N=329). In the presence of Quis (teal), the same two FRET peaks are seen except with different relative proportions of the two states. (e) The addition of G_q_ to the Quis-alone sample shifts the population to the high FRET states, Intermediate 3b (~ 0.75) and Fully Active (~ 0.9) at the expense of the Intermediate 2a (~ 0.6) and Intermediate 1a (~ 0.25) peaks. For the CDPPB alone sample, the addition of G_q_ results in the appearance of a high FRET peak with a decrease, but not complete disappearance of the Intermediate 2b (~ 0.6) and Intermediate 1b (~ 0.25) peaks. (f) 3Dflex analysis of frames 0 and 40 showing a change in distance between the CRDs. (g) Example smFRET traces showing donor (green), and acceptor (red) intensity values as well as the calculated FRET values (blue) for a series of ligand conditions with and without G_q_.

## References

1. Pin, J.-P. & Bettler, B. Organization and functions of mGlu and GABAB receptor complexes. Nature 540, 60–68 (2016).

2. Koehl, A. et al. Structural insights into the activation of metabotropic glutamate receptors. Nature 566, 79–84 (2019).

3. Moustaine El, D., et al. Distinct roles of metabotropic glutamate receptor dimerization in agonist activation and G-protein coupling. Proc. Natl. Acad. Sci. U.S.A. 109, 16342–16347 (2012).

4. Gasparini, F. & Spooren, W. Allosteric modulators for mGlu receptors. Curr Neuropharmacol 5, 187–194 (2007).

5. Ritchie, T. K. et al. Chapter 11 - Reconstitution of membrane proteins in phospholipid bilayer nanodiscs. Methods Enzymol 464, 211–231 (2009).

6. Seven, A. B. et al. G-protein activation by a metabotropic glutamate receptor. Nature 595, 450–454 (2021).

7. Nasrallah, C. et al. Agonists and allosteric modulators promote signaling from different metabotropic glutamate receptor 5 conformations. Cell Rep 36, 109648 (2021).

8. Doré, A. S. et al. Structure of class C GPCR metabotropic glutamate receptor 5 transmembrane domain. Nature 511, 557–562 (2014).

9. Christopher, J. A., Doré, A. S. & Tehan, B. G. Potential for the Rational Design of Allosteric Modulators of Class C GPCRs. Curr Top Med Chem 17, 71–78 (2017).

10. Christopher, J. A. et al. Structure-Based Optimization Strategies for G Protein-Coupled Receptor (GPCR) Allosteric Modulators: A Case Study from Analyses of New Metabotropic Glutamate Receptor 5 (mGlu5) X-ray Structures. J Med Chem 62, 207–222 (2019).

11. Hlavackova, V. et al. Evidence for a single heptahelical domain being turned on upon activation of a dimeric GPCR. EMBO J 24, 499–509 (2005).

12. Fang, W. et al. Structural basis of the activation of metabotropic glutamate receptor 3. Cell Res 32, 695–698 (2022).

13. Isberg, V. et al. Generic GPCR residue numbers - aligning topology maps while minding the gaps. Trends Pharmacol. Sci. 36, 22–31 (2015).

14. Punjani, A. & Fleet, D. J. 3D variability analysis: Resolving continuous flexibility and discrete heterogeneity from single particle cryo-EM. J. Struct. Biol. 213, 107702 (2021).

15. Punjani, A. & Fleet, D. J. 3DFlex: determining structure and motion of flexible proteins from cryo-EM. Nature Methods 2016 14:1 (2023). doi:10.1038/s41592-023-01853-8

16. Lin, S. et al. Structures of Gi-bound metabotropic glutamate receptors mGlu2 and mGlu4. Nature 594, 583–588 (2021).

17. Mansoor, S. E., Dewitt, M. A. & Farrens, D. L. Distance mapping in proteins using fluorescence spectroscopy: the tryptophan-induced quenching (TrIQ) method. Biochemistry 49, 9722–9731 (2010).

18. Liauw, B. W.-H., Afsari, H. S. & Vafabakhsh, R. Conformational rearrangement during activation of a metabotropic glutamate receptor. Nat Chem Biol 17, 291–297 (2021).

19. Liauw, B. W.-H. et al. Conformational fingerprinting of allosteric modulators in metabotropic glutamate receptor 2. Elife 11, (2022).

20. Vafabakhsh, R., Levitz, J. & Isacoff, E. Y. Conformational dynamics of a class C G-protein-coupled receptor. Nature 524, 497–501 (2015).

21. Gregorio, G. G. et al. Single-molecule analysis of ligand efficacy in β2AR-G-protein activation. Nature 547, 68–73 (2017).

22. Punjani, A., Rubinstein, J. L., Fleet, D. J. & Brubaker, M. A. cryoSPARC: algorithms for rapid unsupervised cryo-EM structure determination. Nature Methods 2016 14:1 14, 290–296 (2017).

23. Punjani, A., Zhang, H. & Fleet, D. J. Non-uniform refinement: adaptive regularization improves single-particle cryo-EM reconstruction. Nature Methods 2016 14:1 17, 1214–1221 (2020).

24. Pettersen, E. F. et al. UCSF Chimera--a visualization system for exploratory research and analysis. J Comput Chem 25, 1605–1612 (2004).

25. Adams, P. D. et al. The Phenix software for automated determination of macromolecular structures. Methods 55, 94–106 (2011).

26. Terwilliger, T. C., Sobolev, O. V., Afonine, P. V. & Adams, P. D. Automated map sharpening by maximization of detail and connectivity. Acta Crystallogr D Struct Biol 74, 545–559 (2018).

27. Tan, Y. Z. et al. Addressing preferred specimen orientation in single-particle cryo-EM through tilting. Nature Methods 2016 14:1 14, 793–796 (2017).

28. Emsley, P., Lohkamp, B., Scott, W. G. & Cowtan, K. Features and development of Coot. Acta Crystallogr. D Biol. Crystallogr. 66, 486–501 (2010).

29. Juette, M. F. et al. Single-molecule imaging of non-equilibrium molecular ensembles on the millisecond timescale. Nature Methods 2016 14:1 13, 341–344 (2016).

30. Haran, G. Noise reduction in single-molecule fluorescence trajectories of folding proteins. Chemical Physics 307, 137–145 (2004).

