## Extended Data Table 1 for "Step-wise activation of a Family C GPCR"

### Cryo-EM data collection, refinement and validation statistics

| | CDPPB bound inactive mGlu5 (EMDB-41069) (PDB 8T6J) | Quis bound intermediate mGlu5-Nb43 (EMDB-41092) (PDB 8T7H) | Quis bound active mGlu5-Nb43 (EMDB-41099) (PDB 8T8M) | Quis+CDPPB bound active mGlu5-Nb43 (EMDB-41139) (PDB 8TAO) |
|---|---|---|---|---|
| **Data collection and processing** | | | | |
| Magnification | 107000 | 82000 | 82000 | 82000 |
| Voltage (kV) | 300 | 300 | 300 | 300 |
| Electron exposure (e–/Å$^2$) | 52.2 | 50 | 50 | 51.6 |
| Defocus range (μm) | 0.7-2 | 0.7-2 | 0.7-2 | 0.7-2 |
| Pixel size (Å) | 0.8521 | 1.111 | 1.111 | 1.111 |
| Symmetry imposed | C2 | C2 | C2 | C1 |
| Initial particle images (no.) | 3,565,734 | 4,317,126 | 4,317,126 | 5,033,648 |
| Final particle images (no.) | 246,232 | 211,019 | 183,295 | 245,439 |
| Map resolution (Å) | 3.5 | 3.3 | 3.0 | 2.9 |
| FSC threshold | 0.143 | 0.143 | 0.143 | 0.143 |
| Map resolution range (Å) | 2.8-5.6 | 3.0-7.0 | 3.0-7.0 | 2.5-7.2 |
| **Refinement** | | | | |
| Initial model used (PDB code) | 6N52 | 6N51 and 6N52 | 6N51 | 6N51 |
| Model resolution (Å) | 3.4 | 3.5 | 3.2 | 3.1 |
| FSC threshold | 0.5 | 0.5 | 0.5 | 0.5 |
| Model resolution range (Å) | 31.6-3.4 | 38.5-3.5 | 42.6-3.2 | 2.9-3.1 |
| Map sharpening *B* factor (Å$^2$) | -22.85 | -59.41 | -25.33 | -51.06 |
| Model composition | | | | |
| Non-hydrogen atoms | 11652 | 13100 | 13146 | 13422 |
| Protein residues | 1534 | 1794 | 1770 | 1794 |
| Ligands | 2 | 2 | 2 | 3 |
| *B* factors (Å$^2$) | | | | |
| Protein | 60.67 | 30.43 | 36.58 | 45.08 |
| Ligand | 55.65 | 17.07 | 11.77 | 46.72 |
| R.m.s. deviations | | | | |
| Bond lengths (Å) | 0.002 | 0.003 | 0.003 | 0.002 |
| Bond angles (°) | 0.579 | 0.597 | 0.534 | 0.509 |
| Validation | | | | |
| MolProbity score | 1.69 | 1.73 | 1.64 | 1.48 |
| Clashscore | 6.09 | 7.94 | 6.10 | 4.70 |
| Poor rotamers (%) | 0.00 | 0.00 | 0.00 | 0.00 |
| Ramachandran plot | | | | |
| Favored (%) | 94.88 | 95.73 | 95.66 | 96.40 |
| Allowed (%) | 5.12 | 4.27 | 4.34 | 3.60 |
| Disallowed (%) | 0.00 | 0.00 | 0.00 | 0.00 |
